# Comprehensive comparison of female germ cell development *in vitro* and *in vivo* identifies epigenetic gene regulation crucial for oocyte development and embryonic competence

**DOI:** 10.1101/2023.02.15.528760

**Authors:** Eishi Aizawa, Evgeniy A. Ozonov, Yumiko K. Kawamura, Charles-Etienne Dumeau, So Nagaoka, Mitinori Saitou, Antoine H. F. M. Peters, Anton Wutz

## Abstract

Germ cells are the origin of new individuals. Hence, specifying germ cell identity is crucial for reproduction. The recent establishment of *in vitro* culture systems for generating oocytes from mouse pluripotent stem cells provides a basis for progress in studies of oogenesis and reproductive technology. However, currently the developmental competence of *in vitro* generated oocytes is low compared to *in vivo* grown oocytes. The causes underlying poor oocyte quality remain to be determined. By reconstituting germ cell development in culture from different developmental starting points within gametogenesis, we show that the differentiation of primordial germ cells (PGCs) and primordial germ cell-like cells (PGCLCs) to growing oocytes (GROs), as well as the subsequent growth of follicles are critical culture steps for specifying competence of fully-grown oocytes (FGOs) for preimplantation development. A systematic comparison of transcriptomes of single oocytes having undergone different *in vitro* culture trajectories identifies genes normally upregulated during oocyte growth to be susceptible for mis-regulation during *in vitro* oogenesis. Many of such genes have been described as targets of Polycomb repressive complexes (PRCs). Deregulation of Polycomb repression therefore likely perturbs the accumulation of cytoplasmic factors and/or setting of chromatin states in FGOs that are required for embryonic development after fertilization. Conversely, *in vitro* derived oocytes often displayed failure of zygotic genome activation (ZGA) and abnormal acquisition of 5-hydroxymethylcytosine (5hmC) on maternal chromosomes after activation. In addition, subcellular delocalization of pyruvate dehydrogenase (PDH) and of STELLA were observed suggesting new molecular markers for defective oocyte development. Our study identifies epigenetic regulation at an early stage of oogenesis as crucial for developmental competence and suggests specific *in vitro* culture steps as targets for improving oocyte quality.

**Highlights:**

- Single cell transcriptomics and functional assessment of oocyte development from pluripotent stem cells in culture in a stage-specific manner provides a comprehensive resource for comparisons to oogenesis *in vivo*.
- Culture steps for growth and differentiation of reconstituted follicles are critical for defining embryonic competence of *in vitro* generated oocytes.
- Zygotic genome activation failure and epigenetic impairment are hallmarks of i*n vitro*-generated oocytes that fail to develop after activation or fertilization.
- Computational analysis of gene expression changes and chromatin modification patterns identifies specific gene sets that indicate that Polycomb mediated repression is vulnerable during *in vitro* folliculogenesis.

## Introduction

In animals with sexual reproduction, germ cells are the source of totipotent cells, from which new individuals can develop. Whereas oocytes and spermatozoa transmit their genomes and epigenetic information to the offspring, the oocyte also provides cytoplasmic components that are crucial for development of the embryo after fertilization. Mutations in germ cells are inherited by the offspring and drive genetic variation in species, and can cause embryonic lethality or disorders (Ellegren and Galtier, 2016). How gametes develop to facilitate a totipotent configuration after fertilization remains to be elucidated. In mammals, studying the female germline is challenging as only a small number of germ cells develops to mature oocytes. In addition, tracing germ cell development in the embryo is difficult. For overcoming experimental limitations, *in vitro* culture systems for developing oocytes have been considered for over half a century (Odor and Blandau, 1971). At the beginning of the 2000s, several studies reported the generation of germ cells and mature gametes from pluripotent stem cells (PSCs) including embryonic stem cells (ESCs), induced pluripotent stem cells (iPSCs), epiblast stem cells and embryonic germ cells (Eguizabal et al., 2009; Nayernia et al., 2006; Ohinata et al., 2009; Qing et al., 2007; Toyooka et al., 2003). Especially, two studies succeeded in generating primordial germ cell-like cells (PGCLCs), which gave rise to functional spermatozoa and oocytes, from mouse ESCs and iPSCs by 2-step culture using a cocktail of growth factors (Hayashi et al., 2012; Hayashi et al., 2011). Subsequently, Hikabe and coworkers reported the complete development of female germ cells in culture, thereby enabling the generation of mature metaphase II (MII) oocytes from mouse PSCs including ESCs and iPSCs (Hikabe et al., 2016). This important advance has been recently applied to studying mechanisms of female germ cell development including the dormant state in primordial follicles, effects of sex chromosomes, and transcription factors (TFs) involved in oocyte growth (Hamada et al., 2020; Hamazaki et al., 2021; Nagamatsu et al., 2019; Shimamoto et al., 2019). It has also been used to study the kinetics and efficiency of X-chromosome inactivation and reactivation in female germ cells (Severino et al., 2022). However, oocytes developed *in vitro* have variable potential for embryogenesis. The success rate of full-term development from 2-cell embryos generated from *in vitro* derived MII oocytes is substantially lower (0.9%, 26/2,753) than that of embryos generated using oocytes from superovulated mice (61.7%, 37/60) (Hikabe *et al*., 2016).

During gametogenesis, germ cells undergo extensive epigenetic reprogramming. Following the specification of PGCs and during their migration to gonads between embryonic day (E) 6.5 and E13.5, global CpG methylation levels rapidly decrease (Seisenberger et al., 2012). In parallel, global changes of histone modifications occur. In particular, reduced histone H3 lysine 9 dimethylation (H3K9me2) and elevated histone H3 lysine 27 trimethylation (H3K27me3) are associated with the PGC genome (Hajkova et al., 2008; Seki et al., 2005; Seki et al., 2007). Female PGCs enter meiosis around E13.5 and maintain their DNA largely devoid of methylation. Shortly after birth, primordial follicles emerge, which contain oocytes in meiotic arrest until ovulation (Shirane et al., 2013; Smallwood et al., 2011). The subsequent establishment of proper DNA methylation in the oocyte genome is important for controlling imprinted expression during embryogenesis (Kaneda et al., 2004). Allelic DNA methylation established at imprinting control regions in gametes regulates parental allele-specific expression of imprinted genes in embryos (Tucci et al., 2019). After primordial follicles exit the dormant state, *de novo* DNA methylation is established in growing oocytes (GROs) by the *de novo* DNA methyltransferases, DNMT3A and DNMT3L, in a dynamic interplay with opposing histone methylation pathways (Stäubli and Peters, 2021; Tucci *et al*., 2019). While histone H3 lysine 36 dimethylation and trimethylation (H3K36me2/me3) recruits DNMT3A/3L to chromatin, histone H3 lysine 4 dimethylation and trimethylation (H3K4me2/me3) inhibits DNMT3A/3L catalytic function in oocytes (Ciccone et al., 2009; Ooi et al., 2007; Stewart et al., 2015; Zhang et al., 2010). Polycomb group proteins also contribute to defining developmental competence by silencing differentiation-inducing genes and mediating spatial interactions between genome regions that are marked by H3K27me3 (Du et al., 2020; Posfai et al., 2012). Polycomb group proteins are observed in two major chromatin modifying Polycomb Repressive Complexes, PRC1 and PRC2, which catalyze mono-ubiquitination of histone H2A at lysine 119 (H2AK119ub1) and H3K27me3, respectively (Blackledge and Klose, 2021). Recently, marking of broad genomic regions with H3K27me3 in oocyte genomes was identified to mediate paternal X-chromosome inactivation as well as non-canonical imprinting, causing maternal allele-specific repression of dozens of genes in preimplantation embryos and extraembryonic placental tissues (Chen et al., 2019; Hanna et al., 2019; Inoue et al., 2017a; Inoue et al., 2017b). In GROs, PRC1 functions upstream of PRC2 to define maternal H3K27me3-dependent imprints (Mei et al., 2021). Establishing the proper chromatin configuration during oocyte growth is, thus, a crucial factor for oocyte quality and developmental competence.

Here, we perform a detailed comparison between oocyte development *in vitro* and *in vivo* for identifying potential causes that impair the integrity of oocytes during culture. We first recapitulate oocyte development from PSCs *in vitro* with overall similar rates as previous studies (Hikabe *et al*., 2016). We then compare *in vitro* oocyte development from different developmental starting points of gametogenesis to define critical culture steps. Our data show that the differentiation from PGCs and PGCLCs to GROs, as well as the subsequent growth of follicles are critical for specifying competence of fully-grown oocytes (FGOs) for preimplantation development. Developmental failure of a large fraction of preimplantation embryos from *in vitro*-derived oocytes can be explained by failure of zygotic genome activation (ZGA) and abnormal acquisition of 5-hydroxymethylcytosine (5hmC), which further correlated with inactive pyruvate dehydrogenase (PDH) and the mislocalization of STELLA in the cytoplasm, respectively. Comprehensive transcriptome analysis of individual *in vitro* culture-derived versus *in vivo* generated oocytes identified frequent transcriptional deregulation of genes that are normally repressed by Polycomb group proteins as new molecular factors that are misregulated in *in vitro* generated oocytes. Our study emphasizes epigenetic regulation at an early step of oocyte differentiation as crucial for successful preimplantation development and identifies specific culture steps for attempts of improvement.

## Results

### *In vitro* culture facilitates full female germ cell development from PSCs

We used mouse ESC and iPSC lines that carried Blimp1-Venus and Stella-ECFP reporters (BVSC-ESC and BVSC-iPSC lines) (Hayashi *et al*., 2012; Hikabe *et al*., 2016; Ohinata et al., 2008), as well as an ESC line with a ubiquitously expressed CAG-EGFP reporter (GFP-ESC line) to monitor *in vitro* development of PSCs to mature MII oocytes following a previous report (Hikabe *et al*., 2016). This protocol comprises 4 developmental steps over a span of 45 days. Starting from PSCs, we performed successive *in vitro* PGC differentiation (IVP), *in vitro* oocyte differentiation (IVD), *in vitro* growth (IVG), and *in vitro* maturation (IVM) of oocytes (Figure 1A).

**Figure 1.**
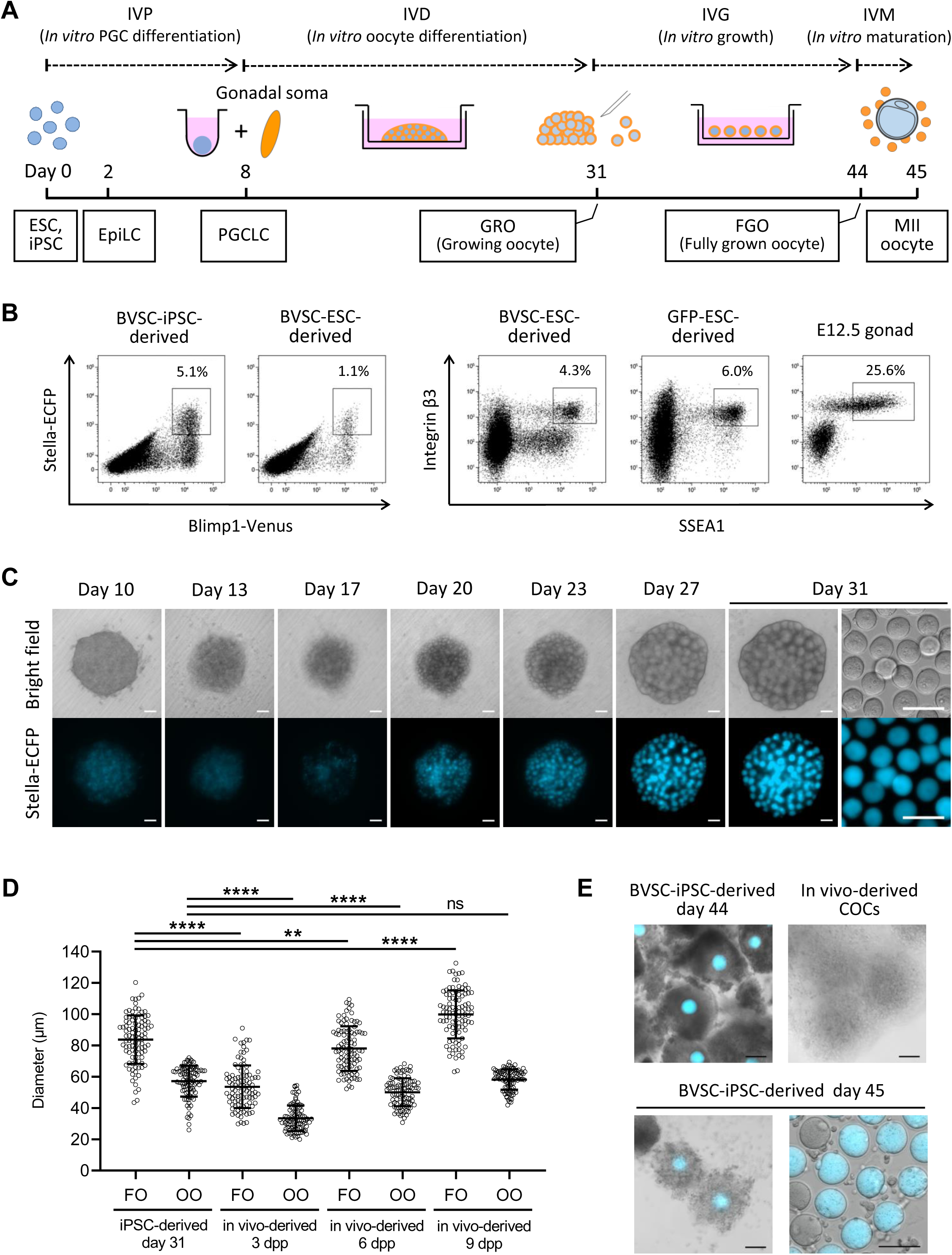
Development of MII oocytes from iPSCs by *in vitro* culture. (A) A schematic illustration of the *in vitro* culture system for the entire development of mouse female germ cells. Developmental stages (dashed line), the culture period (solid line) and developing cell types (square) are shown. (B) A representative flow cytometry analysis of embryoid bodies derived from BVSC-ESC, BVSC-iPSC and GFP-ESC lines at day 8 of the culture. Female E12.5 gonads were analyzed as a control. (C) Development of a BVSC-iPSC-derived rOvary from day 10 to 31 in culture. At day 31, oocytes were harvested from the rOvary (right). Scale bar, 100 µm. (D) Oocyte and follicle diameter of iPSC-derived follicles at day 31, and *in vivo*-derived follicles at 3, 6 and 9 dpp. FO, follicle; OO, oocyte. **** P < 0.0001; ** P < 0.01; ns, non-significant. (E) IVM of iPSC-derived-follicles. Follicles at day 44 before IVM (left top), expanded follicles at day 45 after IVM (left bottom) and collected MII oocytes at day 45 (right bottom) are shown by merging bright field images with Stella-ECFP expression. Bright field image of *in vivo*-derived COCs collected after superovulation of a mouse is shown as control. COCs, cumulus-oocyte complexes. Scale bar, 100 µm.

During 8 days of IVP PSCs differentiate first into epiblast-like cells (EpiLCs) for two days and subsequently into PGCLCs. In BVSC-ESC and BVSC-iPSC lines expression of Blimp1-Venus and Stella-ECFP was observed from day 4 and 6, respectively (Figure S1A). Flow cytometry analysis showed that 1.1% and 5.1% of cells in day 8 embryoid bodies (EBs) generated from BVSC-ESCs and BVSC-iPSCs, respectively, were double positive for both Blimp1-Venus and Stella-ECFP (Figure 1B). Immunostaining for SSEA1 and integrin β3, two surface markers for PGCLCs (Hayashi *et al*., 2011), revealed 4.3% and 6.0% double positive cells in BVSC-ESC and GFP-ESC derived EBs, further confirming the induction of PGC cell fate. Oogenesis requires interactive signals between oocytes and the surrounding gonadal somatic cells, which results in the formation of follicles (Frost et al., 2021; O’Connell and Pepling, 2021). IVD mimics *in vivo* development, by aggregating PGCLCs with somatic cells isolated from E12.5 female gonads in low-binding plates. We prepared such reconstituted ovaries (rOvaries) and cultured them on membranes of transwell plates for 21 days, resulting in the emergence of oocytes derived from BVSC-ESCs and BVSC-iPSCs at day 31 of the culture (Figure 1C, S1B). Some oocytes were observed that lacked Stella-ECFP expression. These oocytes likely originated from incomplete depletion of germ cells from the gonadal somatic cells that we used to form rOvaries, as has been observed previously (Hikabe *et al*., 2016; Yoshino et al., 2021).

To estimate the developmental stage of oocytes and follicles, we compared the diameters of BVSC-iPSC-derived oocytes and follicles to *in vivo* grown oocytes and follicles (Figure 1D, S2A). These measurements showed that the size of iPSC-derived oocytes at day 31 (mean, 57.1 µm) was closest to those of 9 days postpartum (dpp) oocytes (mean, 58.2 µm). The size of iPSC-derived follicles (mean, 83.7 µm) was comparatively close to the size of 6 dpp follicles (mean, 78.0 µm). These results indicate that most iPSC-derived oocytes at day 31 correspond to GROs in primary and secondary follicles that are prevalent in prepubertal ovaries. The data further suggest a reduced rate of proliferation and/or growth of granulosa cells surrounding the GRO during IVD.

To enable further development of primary and secondary follicles to antral and preovulatory stages following the IVG protocol, we mechanically separated follicles in rOvaries at day 31 as described previously (Hikabe *et al*., 2016). Separation of some of the *in vitro*-derived follicles in rOvaries caused denudation of GROs from follicles indicating a fragile follicular structure. In contrast, denudation rarely occurred during dissection of follicles from 6 or 9 dpp ovaries. To overcome the problem of denudation, we tested 9 batches of commercial fetal bovine serum (FBS) and a serum replacement for IVG culture (Figure S3A) and performed isolation of single follicles from rOvaries. Our data show a strong influence of the serum on frequencies of denudation of GROs (Figure S3B). We identified a commercial FBS (Life Technologies, A3161001), which enabled efficient isolation of intact follicles (79.2%) for successive experiments. We further evaluated dissection of rOvaries into clusters of either 1-3 or 4-10 follicles for IVG (Figure S2B, S2C). At day 44 of IVG, each follicle was categorized into 3 groups according to its diameter (0-200 µm; 200-400 µm; over 400 µm) by measuring the longest part in a follicle under a stereomicroscope. The development of follicles was consistent with reports that showed diameters of some follicles reached over 400 µm after IVG (Hikabe *et al*., 2016; Morohaku et al., 2017; Morohaku et al., 2016). We observed a higher proportion of follicles with their diameters over 400 µm in experiments using larger groups of 4-10 follicles (19.6%) compared to 1-3 follicles (12.0%).

At day 44 of the culture, follicles with diameters of greater than 200 µm were harvested and subjected to IVM. After IVM, oocytes and surrounding somatic cells formed expanded cumulus-oocyte complexes (COCs) with a similar morphology to *in vivo*-derived COCs (Figure 1E). Oocytes with first polar bodies, representing MII oocytes, were observed in some COCs derived from BVSC-ESCs and BVSC-iPSCs (Figure 1E, S1C). Overall, we could recapitulate *in vitro* oogenesis with similar efficiency as previous reports.

### Abnormalities of rOvaries and oocytes associated with *in vitro* development

In some rOvaries, we encountered abnormal development of oocytes and follicles during IVD (Table S1; Figure S3C). Small cells protruded from rOvaries in about 16% of samples when we used PGCLCs expressing Blimp1-Venus and Stella-ECFP, and overgrew the culture. This effect was even more pronounced, when we sorted PGCLCs that were derived from GFP-ESC and BVSC-ESC lines for SSEA1 and integrin β3 expression. The observation that all rOvaries were overgrown by small cells suggests that sorting for SSEA1 and integrin β3 expression did not sufficiently enrich for PGCLCs competent for follicle formation.

Following IVM, most of PSC-derived MII oocytes at day 45 of *in vitro* culture were almost indistinguishable from *in vivo* grown MII oocytes (Figure 1E, S1C). In 7% of PSC-derived oocytes (75/1,043) at day 45 we observed small cells on the inner side of the zona pellucida (Figure S3E, F), and some oocytes had a split zona pellucida into 2 branches that resulted in embryos with contaminating cells (Figure S3F). In contrast to the overgrowth of rOvaries, the contaminating cells within and abnormal morphology of the zona pellucida did not affect follicular development.

### IVD and IVG critically define embryonic competence of oocytes

We next assessed the competence of BVSC-iPSC-derived COCs for preimplantation development after IVM and fertilization with sperm (Figure 2A-B; Table 1). After IVF, zygotes were observed that progressed through cleavage divisions and developed into blastocysts at the ratio of 1.7% (14/838). We also parthenogenetically activated BVSC-iPSC-derived oocytes at day 45 to measure their developmental competence without factors from spermatozoa (Figure S2D; Table 1). Using both methods, we found that iPSC-derived oocytes exhibited notably lower developmental rates at all stages of preimplantation development than control *in vivo*-derived oocytes.

**Figure 2.**
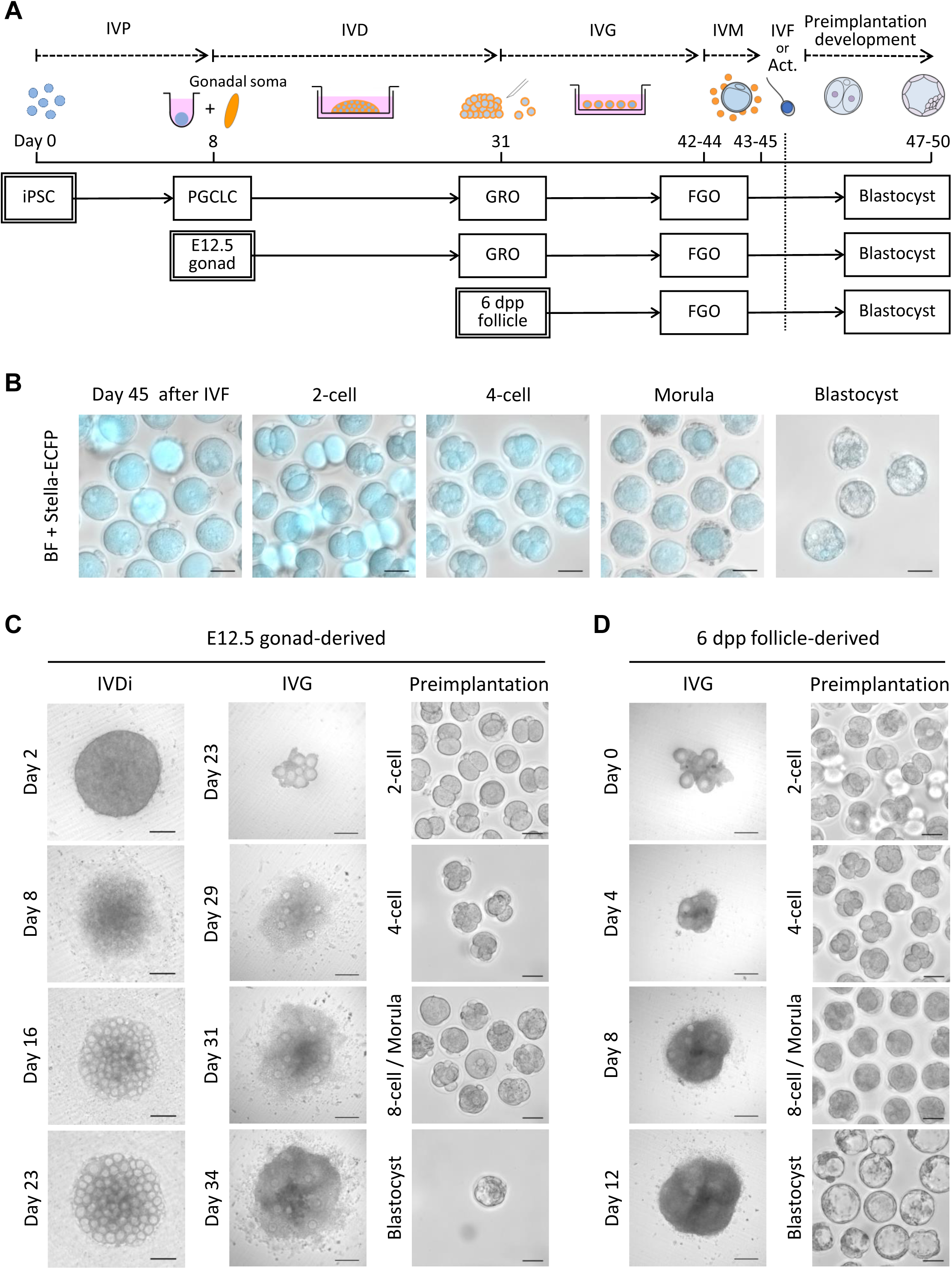
Comparison of *in vitro* development of iPSC-, E12.5 gonad- and 6 dpp follicle-derived oocytes and embryos. (A) A scheme to assess developmental stages of the *in vitro* culture. A similar culture protocol was applied to 3 different cell types (iPSCs, E12.5 gonads, and 6 dpp follicles), starting from each developmental stage (germ cell differentiation, IVD and IVG). After IVM, oocytes derived from each cell type were subjected to IVF or parthenogenetic activation, and preimplantation development was followed *in vitro*. Act, Activation. (B) Preimplantation development of BVSC-iPSC-derived oocytes after IVF. Stella-ECFP expression was merged with bright field images. Scale bar, 50 µm. (C) Development of E12.5 gonad-derived rOvary, follicles and embryos after IVF. Scale bar, 200 µm (IVD and IVG) and 50 µm (preimplantation development). (D) Development of 6 dpp follicles and 6 dpp follicle-derived embryos after IVF. Scale bar, 200 µm (IVG) and 50 µm (preimplantation development).

**Table 1.** Summary of preimplantation development of *in vitro*- and *in vivo*-derived oocytes

| Source of oocytes | IVF or Activation | No. of exp. | No. of rOva | No. of all oocytes | No. of MII oocytes | No. of 2-cell embryo | No. of 4-cell embryo | No. of 8-cell embryo / morulae | No. of blastocysts (% of all oocytes) |
| --- | --- | --- | --- | --- | --- | --- | --- | --- | --- |
| BVSC-iPSC-derived | IVF | 3 | 72 | 838 | Not counted | 208 | 85 | 49 | 14 (1.7%) |
|  | Activation | 18 | 416 | 5701 | 722 | 483 | 43 | 6 | 0 (0%) |
| E12.5 gonad | IVF | 2 | 26 | 584 | Not counted | 121 | 31 | 24 | 5 (0.9%) |
| 6 dpp follicle | IVF | 2 | - | 739 | Not counted | 447 | 297 | 249 | 111 (15.0%) |
| In vivo-derived | IVF | 3 | - | 151 | Not counted | 115 | 104 | 97 | 84 (55.6%) |
|  | Activation | 2 | - | 194 | 187 | 101 | 64 | 50 | 27 (13.9%) |

To identify which *in vitro* culture steps are critical for defining oogenic and embryonic developmental competence we performed a comparative assessment of the different stages of *in vitro* culture versus *in vivo* development (Figure 2A). For this we cultured E12.5 female gonads and 6 dpp follicles in accordance with corresponding *in vitro* culture protocols and compared their developmental competence to those of *in vivo* generated follicles. We chose 6 dpp follicles given that their overall size is comparable to that of PSC-derived follicles after the IVD, even though oocytes from 6 dpp follicles are smaller than those of day 31 PSC-derived oocytes (Figure 1D). This experimental design allowed us to relate developmental efficiency to gene expression profiles of individual oocytes, and to assess the impact of IVP, IVD and IVG on oocyte development by comparing *in vitro* culture and *in vivo* grown germ cells of different developmental stages (Figure 2B, 5A). E12.5 gonads were dissociated to form rOvaries followed by the IVD culture (Figure 2C). E12.5 gonad-derived follicles in rOvaries were dissected at day 23 and subsequently cultured following IVG and IVM. We also dissected follicles from 6 dpp ovaries and cultured them through the IVG and IVM steps (Figure 2D). After IVM, COCs derived from either E12.5 gonads or 6 dpp ovaries were subjected to IVF to assess their competence for preimplantation development.

Comparison of follicle expansion during the IVG demonstrated that about 20% of BVSC-iPSC-derived (19.6%) and E12.5 gonad-derived follicles (22.7%) reached a diameter of over 400 µm. In contrast, double the number of 6 dpp ovary-derived follicles reached a diameter of over 400 µm (48.7%) (Figure 3A). We next monitored preimplantation development and observed development of blastocyst embryos at a rate of 1.7% and 0.9% for BVSC-iPSC-derived and E12.5 gonad-derived oocytes. In contrast, 6 dpp follicle-derived oocytes were 8-fold more likely to develop to blastocysts (15.0%) (Figure 3B; Table 1). Furthermore, E12.5 gonad-derived and BVSC-iPSC-derived oocytes showed significant decreases in the transitions from oocyte to 2-cell embryo and from 2-cell to 4-cell embryo compared to 6 dpp follicle-derived oocytes (Figure 3C-E). These results have two implications. Firstly, rOvaries generated from BVSC-iPSC-derived PGCLCs and E12.5 gonad-derived PGCs possess comparatively similar developmental competences for IVG and preimplantation development. This indicates that PGCLC development *in vitro* is comparable to PGC development *in vivo*, at least for the current IVD settings. Secondly, a substantially reduced level of follicle expansion was observed for BVSC-iPSC-derived and E12.5 gonad-derived follicles compared to 6 dpp ovary-derived follicles during IVG. This result indicates that currently used IVD culture conditions negatively affect oocyte growth and limit embryonic competence. Given the relatively low blastocyst rate of even 6dpp follicle-derived oocytes (15%, Figure 3B), we conclude that conditions for reconstitution and culture of oocyte – granulosa aggregates (IVD) as well as subsequent growth (IVG) require further optimization.

**Figure 3.**
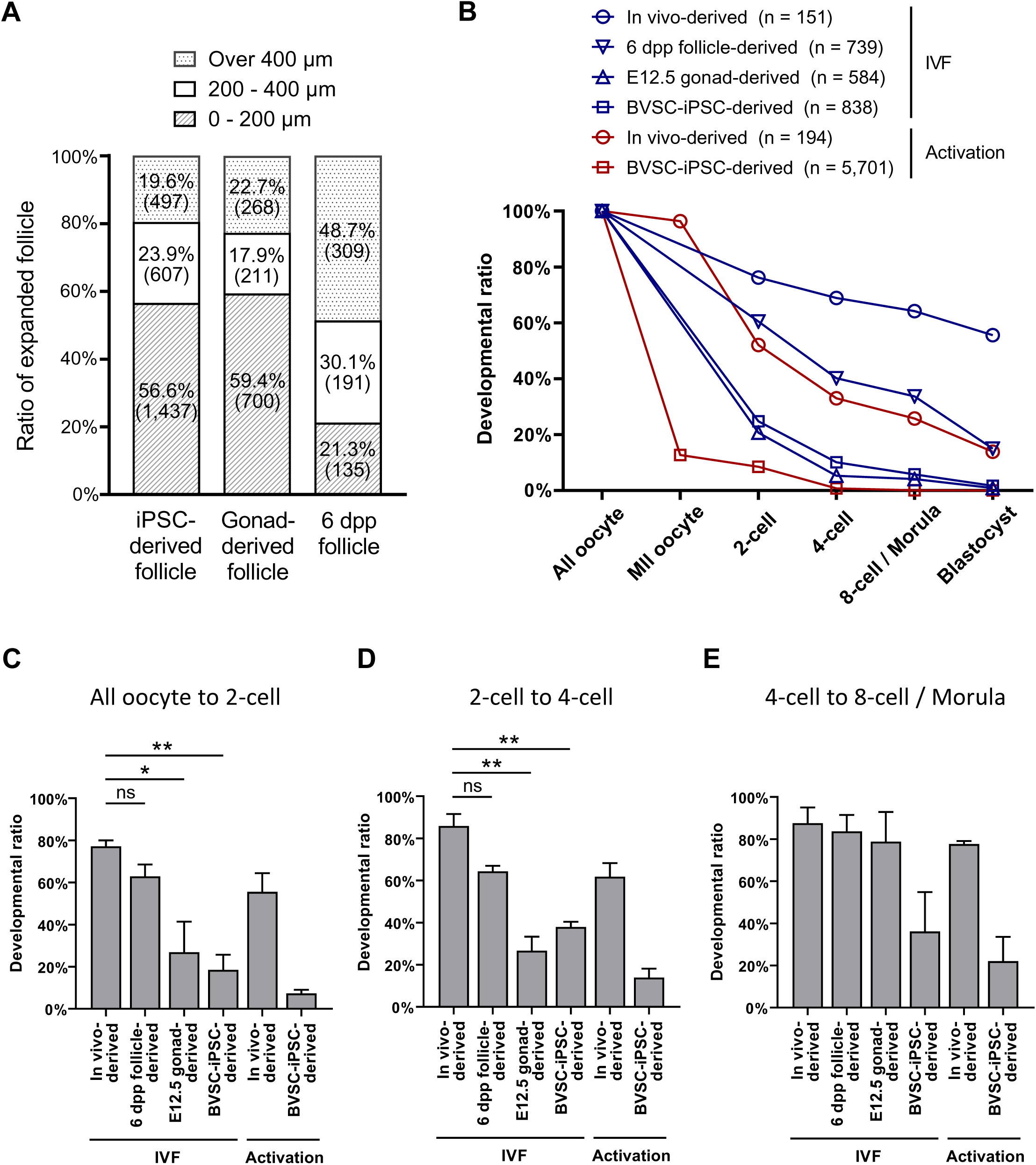
Developmental efficiency of IVG and preimplantation development of PSC-, E12.5 gonad- and 6 dpp follicle-derived oocytes. (A) Summary of follicle development by IVG. The largest diameters of BVSC-iPSC-derived, E12.5 gonad-derived and 6 dpp follicle-derived follicles were measured at day 44, day 34 and day 12 of the culture, respectively. Numbers in brackets represent the number of counted follicles. The data of iPSC-derived follicles is identical to one of the data in Figure S2C. (B) Summary of preimplantation development expressed as a ratio with the number of counted oocytes in each sample set to 100%. Data are from 3 (*in vivo*-derived, IVF), 2 (6 dpp-follicle-derived, IVF), 2 (E12.5 gonad-derived, IVF), 3 (BVSC-iPSC-derived, IVF), 2 (*in vivo*-derived, activation) and 18 (BVSC-iPSC-derived, activation) independent experiments. (C, D and E) Developmental ratio from all oocytes to 2-cell embryos (C), from 2-cell to 4-cell embryos (D), and from 4-cell to 8-cell embryos / morulae (E). N = 3 (*in vivo*-derived, IVF), 2 (6 dpp-follicle-derived, IVF), 2 (E12.5 gonad-derived, IVF), 3 (BVSC-iPSC-derived, IVF), 2 (*in vivo*-derived, activation) and 18 (BVSC-iPSC-derived, activation). * P < 0.05; ** P < 0.01; ns, non-significant.

### Impaired zygotic genome activation and epigenetic regulation is associated with *in vitro* culture of oocytes

The rate of preimplantation development to blastocysts varied widely from 55.6% to 0% among the 6 culture conditions (Figure 3B; Table 1). Parthenogenetic activation and development of haploid embryos from BVSC-iPSC-derived oocytes was the only condition which did not result in blastocyst formation. Although 66.9% of BVSC-iPSC-derived haploid parthenotes developed from MII oocytes to 2-cell embryos (483/722; Table 1), most 2-cell parthenotes notably failed to develop into 4-cell embryos (8.9%, 43/483). This result suggests that BVSC-iPSC-derived oocytes lack or misexpress factors which are critical for the maternal-to-zygotic transition and development beyond the 2-cell stage.

Early embryonic development is initially supported by maternal factors that accumulate in the egg during oogenesis and are progressively replaced by factors expressed in embryos (Zhang et al., 2022). To investigate this point further, we performed immunostaining analysis for key factors of the maternal-to-zygotic transition (Figure 4; Figure S4). To exclude effects from sperm, we studied parthenogenetically activated (PA) 2-cell embryos. We applied a Bromouridine-triphosphate (BrUTP) assay to quantify the extent of nascent transcription. Three quarters of BVSC-iPSC-derived PA 2-cell embryos (18/24) lacked any BrUTP staining indicating that they failed to activate transcription of the embryonic genome (Figure 4A, B). Pyruvate dehydrogenase (PDH) has been reported to serve an important role in ZGA. It localizes in its active non-phosphorylated form in the nucleus of 2-cell embryos (Nagaraj et al., 2017). While all *in vivo* oocyte-derived 2 cell embryos showed nuclear enrichment of PDH, nuclear PDH levels were greatly reduced or even absent in BVSC-iPSC-derived 2-cell embryos (Figure 4C, D). Hence, such abnormal PDH localization may contribute to the observed ZGA failure.

**Figure 4.**
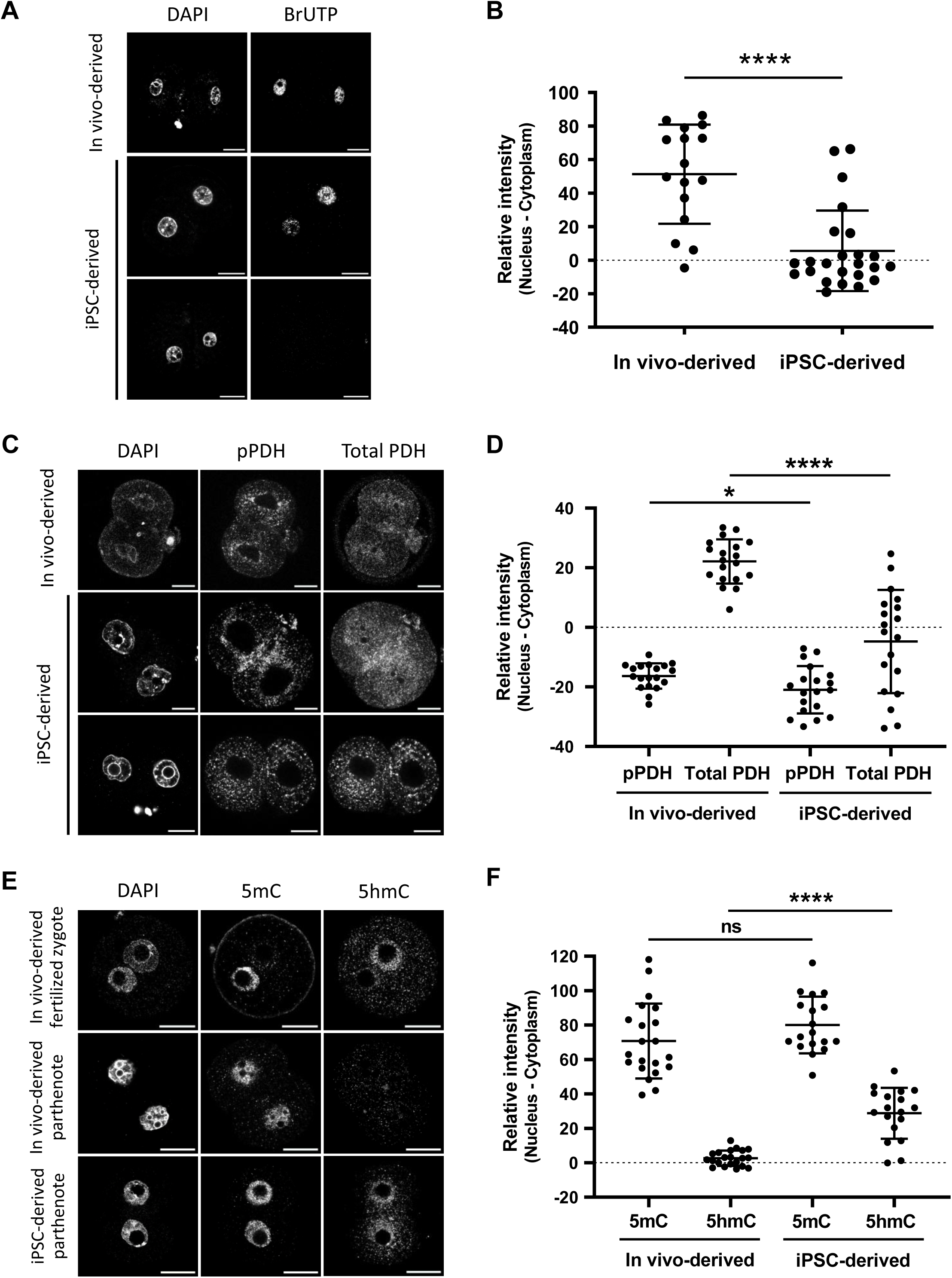
Immunostaining analysis of nascent transcripts, 5mC and PDH in parthenogenetic 2-cell embryos. (A) Representative staining of nascent transcripts after BrUTP incorporation in PA 2-cell embryos. (B) Quantitative data of nascent transcripts after BrUTP incorporation in PA 2-cell embryos. n = 16 (*in vivo*-derived) and n = 24 (iPSC-derived). (C) Representative staining of pPDH and total PDH in PA 2-cell embryos. (D) Quantitative data of pPDH and total PDH staining in PA 2-cell embryos. n = 19 (*in vivo*-derived) and n = 19 (iPSC-derived). (E) Representative staining of 5mC and 5hmC in PA 2-cell embryos. A fertilized zygote is shown as control. (F) Quantitative data of 5mC and 5hmC staining in PA 2-cell embryos. n = 21 (*in vivo*-derived) and n = 18 (iPSC-derived). Scale bar, 20 µm. * P < 0.05; **** P < 0.0001; ns, non-significant.

We next asked if epigenetic marks of maternal chromosomes could provide hints for a potential cause of defects of culture derived oocytes. H3K4me3 and H3K27me3 have been associated with active and repressed genes, respectively, and both modifications are inherited on maternal chromatin from eggs to 2-cell embryos (Stäubli and Peters, 2021). Immunostaining analysis demonstrated a small increase of H3K27me3 levels in BVSC-iPSC-derived PA 2-cell embryos, while levels of H3K4me3 and H3K27 acetylation (H3K27ac), which marks active enhancers (Hanna et al., 2018a), were comparable to those in *in vivo*-derived controls (Figure S4C-F). Furthermore, we observed increased levels of H3K4 acetylation (H3K4ac) in BVSC-iPSC-derived 2-cell embryos, which might negatively regulate the loading of the chromosome passenger complex during metaphase (Niedzialkowska et al., 2022). DNA methylation of the gametic genomes undergoes dramatic changes after fertilization. Whereas oocyte-derived 5-methylcytosine (5mC) undergoes passive demethylation over successive cell divisions in the early embryo, the paternal genome undergoes active DNA demethylation in zygotes, in part through TET-mediated conversion of 5mC to 5hmC (Amouroux et al., 2016; Nakamura et al., 2007; Nakamura et al., 2012). Immunostaining of 5mC and 5hmC in BVSC-iPSC-derived PA 2-cell embryos revealed an unexpected increase in 5hmC levels on maternal chromosomes, while 5mC levels were comparable to those in PA embryos derived from *in vivo* grown oocytes (Figure 4E, F). Binding of the STELLA protein to H3K9me2 on maternal chromosomes has been reported to prevent TET3-mediated oxidation of 5mC into 5hmC (Nakamura *et al*., 2012). Importantly, immunostaining analysis of H3K9me2 and STELLA revealed that some BVSC-iPSC-derived 2-cell embryos exhibited weak nuclear localization of STELLA (Figure S4A, B). Reduced localization of STELLA in the nucleus correlates with enhanced acquisition of 5hmC after activation of *in vitro*-derived oocytes.

### Genes normally up-regulated during oocyte growth are particularly vulnerable for mis-regulation during *in vitro* oogenesis

To identify possible causes underlying the overall low oogenic and embryonic competence of *in vitro*-derived oocytes, we performed extensive comparative transcriptome analysis between *in vivo* and *in vitro* grown germ cells, at different stages of their development. Firstly, we performed RNA-Seq analysis of BVSC-iPSC-derived PGCLCs at day 8 of *in vitro* culture (termed d6PGCLCs, representing PGCLCs 6 days after initiating IVP from EpiLCs) and PGCs isolated from E12.5 female gonads by fluorescence-activated cell sorting (FACS) for both SSEA1 and integrin β3 expression (Figure 1B). Principal component analysis revealed reproducible gene expression differences between d6PGCLCs and E12.5 PGCs which correlated well to previously published RNA-Seq datasets (Ohta et al., 2017; Sasaki et al., 2015) (Figure S5A-C). We identified 773 and 877 up- and down-regulated genes between d6PGCLCs and E12.5 PGCs respectively (Figure S5B; Table S2). Gene ontology analysis revealed that genes down-regulated in PGCLCs serve in various female and male germ cell development related functions, in signaling pathways, in embryonic development and transcriptional processes (Figure S5D, Table S3), hinting towards immature repression by Polycomb Repressive Complexes (PRCs) of such genes (Blackledge and Klose, 2021).

Secondly, we performed RNA-Seq analysis of single oocytes at the GRO and FGO stages to quantify and molecularly dissect the impact of different steps of IVP, IVD and IVG on gene expression. The single cell nature of the experiment also allows to assess the transcriptional heterogeneity among and between *in vitro* and *in vivo* grown oocytes. To do so, we directly isolated GROs and FGOs from mice or produced them in vitro using BVSC-iPSCs, E12.5 gonads, and 6 dpp follicles as sources of starting material (Figure 5A). Among the 162 separately sequenced GRO and FGO oocytes (Table S4), 27% of variance in expression could be assigned to differences between GROs and FGOs. Moreover, 8% of the variance directly relates to differences in *in vitro* culture conditions (Figure 5B). More specifically, expression differences between GROs derived from BVSC-iPSCs (A2) and from E12.5 embryonic gonads (B2) can presumably be attributed to culture differences between *in vitro* PGCLCs versus *in vivo* PGC development, since both cohorts A2 and B2 underwent the same IVD. We refer to such effect as “IVP@GRO” (Figure 5A). Similarly, expression differences between E12.5 gonad-derived GROs (B2), having undergone IVD, versus *in vivo*-derived GROs (D2) can be attributed to the IVD stage (IVD@GRO in Figure 5A). To simplify our model, we assumed that expression differences between iPSC-derived GROs (A2) and *in vivo*-derived GROs (D2) are linear combinations of both effects of IVP and IVD (i.e., IVP@GRO + IVD@GRO). A similar approach was taken to quantify effects resulting from IVP, IVD or IVG and impacting expression in FGOs (@FGO). We utilized a generalized linear model to fit corresponding coefficients for each effect and performed likelihood ratio tests to find genes with statistically significant response to each step of *in vitro* development in GROs and FGOs (with FDR ≤ 5% and | log_2_ (Fold-change) | ≥ 2). Using these criteria, we identified around 100 to 300 genes up- or down-regulated in GROs or FGOs, in response to IVP, IVD or IVG (Figure 5C; Table S5). Gene ontology analysis identified down-regulation of genes involved in Notch signaling and upregulation of genes involved in cell adhesion (e.g. *Cdh12*, *Cdh6*, *Cntn5*, *Dsc3*) resulting from IVP as measured in GROs (IVP@GRO) (Figure S5E; Table S6). The impact of IVP on FGOs resulted in down-regulation of genes involved in Xenobiotic metabolic processes (e.g. *Cypb1b1*, *Cyp2c66*). IVD results in down-regulation of genes in retinoid metabolism in GROs and FGOs (e.g. *Adh1, Aldh1a1*) while genes with roles in cell adhesion, olfactory function and chemokine signaling (*Ccl1*, *Ccl4*, *Cx3cl1*, *Cxcl13*) are overrepresented among upregulated genes in IVD@GRO. IVG impacts on cell adhesion and retinoid metabolism as well, yet in an opposite manner as during IVD, as measured in FGOs (Table S6).

While the cellular impact of altered expression of different genes remains to be explored, we next aimed to understand possible modes of regulation underlying altered gene expression between *in vitro* and *in vivo* development. Therefore, we first clustered all genes into 4 groups according to their dynamics of expression during oocyte growth *in vivo* (Figure 5D). We named these groups LS for genes with low stable expression, UP (up-regulated from GRO to FGO), DN (down-regulated from GRO to FGO), and HS (high stable expression between GRO and FGO). We then investigated which genes in each group were differentially expressed by any of the steps of *in vitro* culture (Figure 5C and D). Among all differentially expressed genes we observed a significant, 2.7-fold over-representation of genes which are normally up-regulated during the growth of GROs to the FGO stage *in vivo* (UP group) (Figure 5E). In addition, we observed that genes with stable high expression both in GROs and FGOs (HS group) were significantly under-represented among affected genes.

To further investigate which gene expression dynamics *in vivo* characterize genes affected by each *in vitro* culture step we performed *χ*^2^ tests for significant over-representation of genes with LS, UP, DN and HS dynamics among up- and down-regulated genes by the modeled effects. Interestingly, we observed that among genes normally up-regulated from GRO to FGO (“UP”), 45 were down-regulated in GROs by IVP, 152 were up-regulated in GROs by IVD, and 71 down-regulated in FGO following IVG (Figure 5F).

Next, we investigated promoter features, such as presence of CpG islands (CGIs) at promoters of genes affected by *in vitro* culture (Sendžikaitė and Kelsey, 2019). Interestingly, we did not observe significant enrichment of either CGI nor non-CGI promoters among affected genes that normally display UP dynamics (Figure 5D and E). In contrast, most genes affected by each stage with LS dynamics were significantly enriched with CGI-driven promoters (Figure S5F). We also performed enrichment analysis of transcription factor motifs within promoters of genes with statistically significant expression response to steps of *in vitro* culture and particular expression dynamics from GRO to FGO *in vivo* (Figure 5F) using R/Bioconductor package *monaLisa* (Machlab et al., 2022) and transcription factor binding profile database *JASPAR2020* (Fornes et al., 2020). Our analysis did, however, reveal only one TF motif with a false discovery rate slightly below our cutoff (FDR ≤ 1%, Figure S5G, Table S7). The RXRA::VDR motif is enriched within promoters of genes which are down-regulated by IVG@FGO and belong to DN group (Figure S5G, Figure 5F) and represents binding specificity of heterodimer between retinoid X receptor alpha (*Rxra*) and vitamin D3 receptor (*Vdr*). However, the *Rxra* gene does not have a significantly altered expression response to IVG@FGO and the *Vdr* gene is not expressed in any of the oocyte cohorts (Table S5), hence it is unlikely that these genes are responsible for observed expression responses of targets to IVG@FGO. Nevertheless, our analysis does not rule out the possibility of other transcription factors with similar sequence specificity to play a role. We finally asked whether any correspondence exists between genes differently expressed in PGCLCs and PGCs and genes affected during *in vitro* oogenesis (Figure 5C, S5B). For genes up- and down-regulated in d6PGCLCs relative to E12.5 PGCs, we did not observe apparent correspondences to the effects of *in vitro* culture in GROs and FGOs (Figure S5H). In summary, our analysis indicates that the observed step-specific effects of *in vitro* culture on gene expression are unlikely explained by either CpG promoter composition, enrichment of TF motifs or by expression differences between PGCLCs and PGCs.

### *In vitro* differentiation results in premature activation of gene expression

We next investigated chromatin modifications at promoters of genes that were affected by the *in vitro* culture. We performed non-parametric Wilcoxon tests for a panel of publicly available ChIP-seq datasets to investigate differences in enrichments for selected chromatin marks between promoters of affected and unaffected genes with the same expression dynamics *in vivo* (groups LS, UP, DN, and HS) and presence or absence of a CGI promoter (Figure 6A). Remarkably, CGI promoters (+/- 1.5 kb around TSS) of affected genes with UP and DN dynamics were enriched with repressive histone marks catalyzed by PRCs, such as H3K27me3 and H2AK119ub1, and depleted for active marks, such as H3K4me3 and H3K27ac, in PGCs, PGCLCs as well as in oocytes including GRO and FGO. This suggests that CGI-driven genes which showed expression changes during normal development from GRO to FGO (groups UP and DN) and were aberrantly expressed in the *in vitro* system are generally controlled by PRCs during oogenesis.

**Figure 5.**
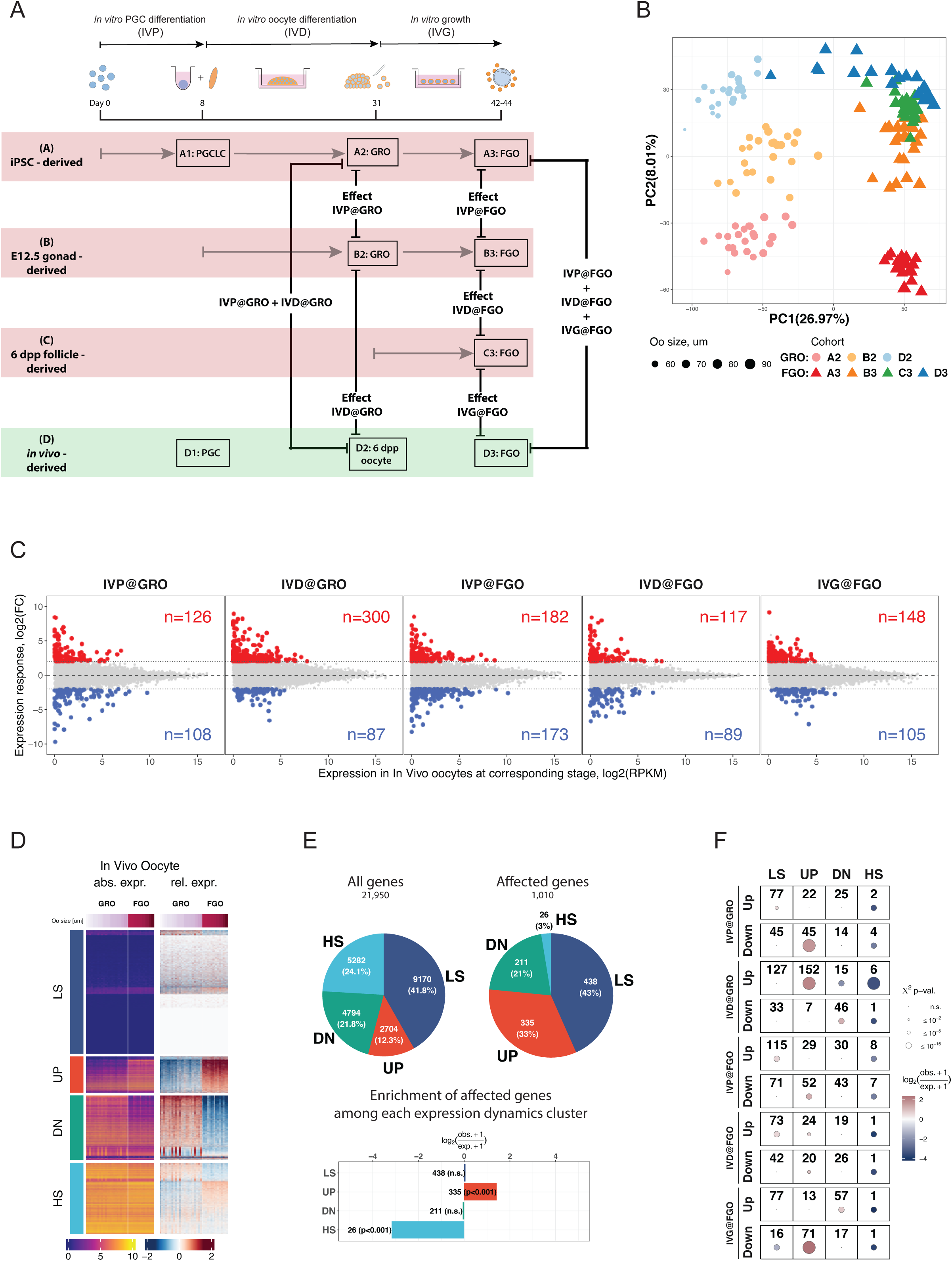
Modelling and quantification of effects of the *in vitro* development on gene expression. (A) Experimental design and illustration of the effects of *in vitro* versus *in vivo* culture used as covariates in a Generalized Linear Model for gene expression. Expression for each gene at corresponding stage (GRO or FGO) is modeled as linear combination of effects of *in vitro* culture stages relative to *in vivo*-derived oocytes. (B) PCA of single oocyte RNA-Seq data used to identify gene expression responses to stages of *in vitro* development. Each point corresponds to a single oocyte scaled, colored and shaped according to corresponding size (um), cohort and developmental stage respectively. (C) Gene expression responses to effects of *in vitro* culture. X-axis represents expression of genes for *in vivo*-derived oocytes at corresponding stage and Y-axis represents the quantified expression responses of each gene (log2(Fold-change)) for each stage. Colored points and numbers in red and blue represent genes whose expression is significantly (with *FDR* ≤ 5% and |*log*_2_(*Fold* – *change*)| ≥ 2) affected by a corresponding stage of the *in vitro* development. (D) Grouping of genes according to the dynamics of expression between GROs and FGOs. All genes were classified into 4 groups, representing genes with Low Stable expression (LS group), up-regulated from GRO to FGO (UP group), down-regulated from GRO to FGO (DN group), and genes with High Stable expression (HS group). (E) Genes which are upregulated in FGOs relative to GROs are overrepresented among genes affected by any of the stages of the *in vitro* culture. Pie chart represents numbers and fraction of genes belonging to each group in (D) among all genes (left pie chart) and genes which are affected in any of the stages of the *in vitro* culture. Results of *χ*^2^-tests and enrichments of each group among affected genes are displayed below the pie charts. (F) Results of *χ*^2^-tests and enrichments of each group among genes with different response (up-regulation or down-regulation) to each stage of the *in vitro* culture. Groups which have *χ*^2^-test p-value larger than 1% are displayed as dots and considered statistically not significant (n.s.).

**Figure 6.**
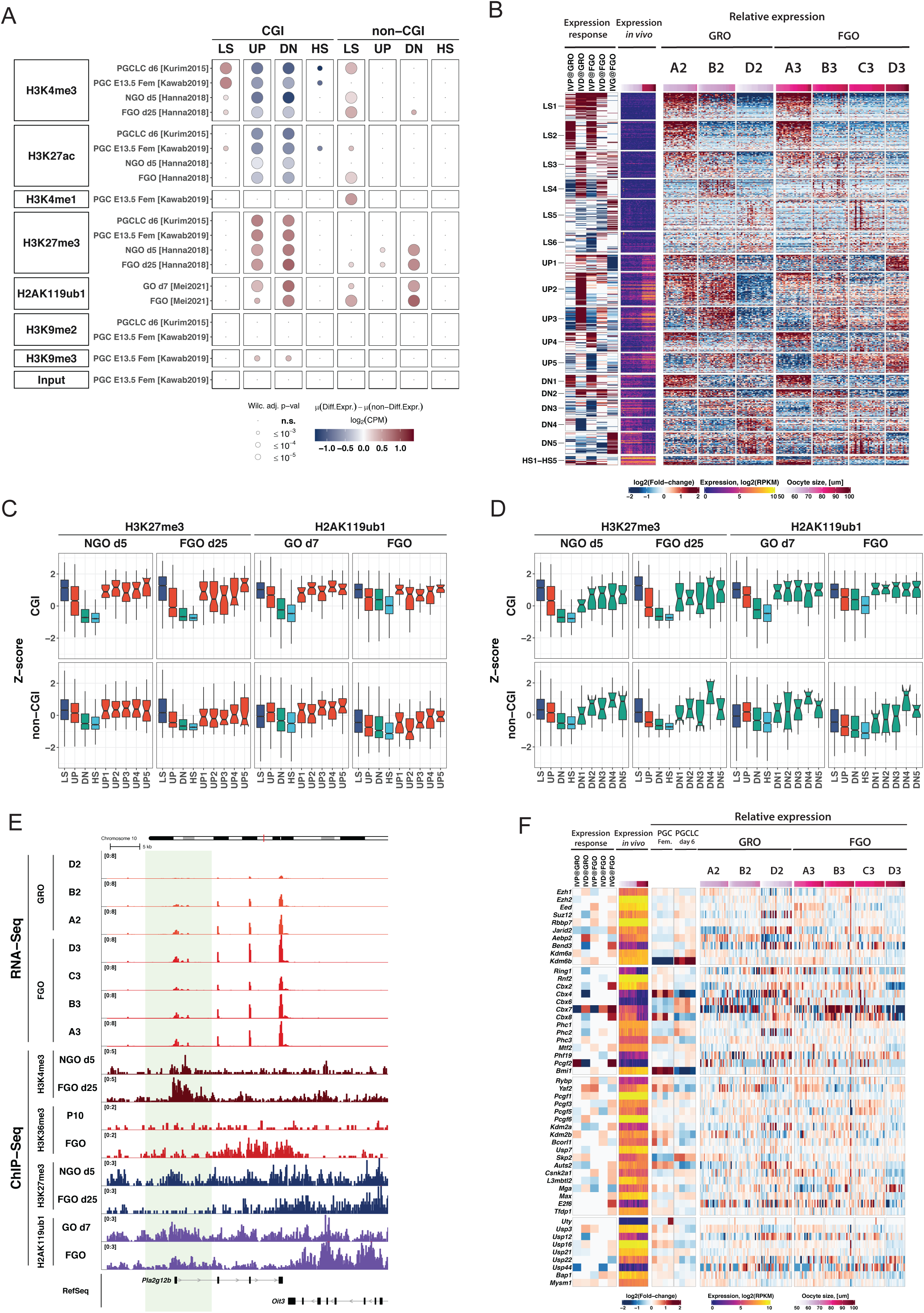
Dynamics of epigenetic chromatin marks during oogenesis is distinct at CGI promoters of genes affected by *in vitro* culture procedure. (A) Differences of histone PTMs in PGC, PGCLC, NGO at day 5 and FGO (data from (Hanna *et al*., 2018b; Kawabata *et al*., 2019; Kurimoto *et al*., 2015; Mei *et al*., 2021)) at promoters of affected genes compared to non-affected genes in the same group. Results of Mann-Whitney tests are displayed for comparison of enrichment of each chromatin mark at promoters of affected genes compared to non-affected genes belonging to the same group. Analyses were separately done for CGI and non-CGI promoter genes. (B) K-means clustering of genes according to combination of gene expression responses to stages of the *in vitro* protocol (heatmap Expression response). Expression in *in vivo* oocytes is shown to illustrate dynamics from GRO to FGO (heatmap Expression *in vivo*). Relative expression of genes in oocytes belonging to different cohorts in GRO (A2, B2, D2) and FGO (A3, B3, C3 and D3) is also depicted. Relative expression for each gene was calculated as the difference between expression (log2(RPKM)) in each oocyte and average expression across all oocytes for GRO and FGO separately. (C) Boxplots displaying enrichments of histone PTMs catalyzed by repressive Polycomb-group complexes at CGI and non-CGI promoters of genes with different expression dynamics during *in vivo* development (LS, UP, DN and HS, see Figure 5C) and affected genes in UP group sub-clusters, defined based on combination of expression responses (as shown in B) (H3K27me3 data from (Hanna *et al*., 2018b), H2AK119ub1 data from (Mei *et al*., 2021)). (D) Boxplots displaying enrichments of histone PTMs catalyzed by repressive Polycomb-group complexes at CGI and non-CGI promoters of genes with different expression dynamics during *in vivo* development (LS, UP, DN and HS, see Fig. 5C) and affected genes in DN group sub-clusters, defined based on combination of expression responses (as shown in B) (H3K27me3 data from (Hanna *et al*., 2018b), H2AK119ub1 data from (Mei *et al*., 2021)). (E) Genomic snapshot of a representative gene *Pla2g12b* belonging to UP2 cluster (see B) and distribution of several chromatin marks. The analysis suggests premature activation of the gene in GRO caused by the IVD of the protocol, possibly due to premature removal of the Polycomb group epigenetic marks H3K27me3 or H2AK119ub1. (F) Expression responses to each stage of *in vitro* development of genes belonging to PRC1 and PRC2 complexes (heatmap *Expression response*, all log2(Fold-change) with *FDR* ≥ 5% are considered non-significant and set to 0). In addition, expression in *in vivo* oocytes (heatmap *Expression in vivo*), and relative expression in PGC and PGCLC, GRO and FGO are displayed (group of heatmaps *Relative expression*). Relative expression for each gene was calculated as difference between expression level (log2(RPKM)) and average expression calculated separately for PGC and PGCLC, GRO, and FGO.

To further elucidate characteristics of genes affected by the *in vitro* development we applied k-means clustering for log-fold-change expression responses (right columns, “relative expression”, Figure 6B) to assign affected genes into groups characterized by combinations of gene expression responses to the *in vitro* culture treatment (left columns, Figure 6B). Interestingly, clusters UP2 and UP3, which showed up-regulated expression from GRO to FGO *in vivo* (UP-group), contained genes whose response to the IVD at GRO was highly positive, indicating aberrant premature activation of these genes during IVD in iPSC- and E12.5 gonad-derived oocytes.

The differences in expression of genes belonging to clusters UP2 and UP3 may possibly be confounded by slight differences in growth of oocytes during their development *in vitro* and *in vivo*, given that oocytes in the A2 and B2 cohorts were larger in size compared to oocytes in the *in vivo* grown D2 cohort (Figure S6A). When taking oocyte diameter into account, our analysis clearly shows that up-regulated expression of genes belonging to UP2 and UP3 in A2 and B2 cohorts was also observed in those oocytes having the same size as oocytes in the D2 cohort (Figure S6B). Moreover, if expression differences observed for UP2 and UP3 genes would have been confounded by differences in oocyte sizes in the three cohorts, we should have observed a general up-regulation of all genes with up-regulation dynamics from GRO to FGO, which we did not (UP group; Figure S6B). In summary, genes in UP2 and UP3 are differently affected by IVD compared to all genes in the UP group. The data suggests that the IVD procedure leads to premature activation of these genes in B2 and partially in A2 cohorts.

### Polycomb-marked genes undergo premature activation during IVD

As we observed a noticeable difference in enrichments of PRC1 and PRC2 chromatin marks at promoters of affected genes belonging to the UP and DN groups, we hypothesized that these genes undergo chromatin remodeling during normal oocyte development. We quantified ChIP-seq enrichments at CGI and non-CGI promoters of genes (+/- 1.5 Kb around TSS) belonging to clusters formed by gene expression dynamics and magnitude of response to *in vitro* differentiation (see Figure 6B, e.g. UP1-UP5, DN1-DN5, etc) and compared them to enrichments of all analyzed genes with different expression dynamics, i.e. LS, UP, DN and HS groups (Figures 6C, D and S6,C-H). As expected, we found that promoters of genes in the LS group which exhibited low stable expression both in GRO and FGO were highly enriched with repressive histone marks, such as H3K27me3 and H2AK119ub1, and showed low enrichments of activating histone marks, such as H3K4me3, both in day 5 non-growing oocytes (NGOs) and FGO (Hanna et al., 2018b) compared to genes in the UP, DN and HS groups (Figure 6C, S6C). In contrast, promoters of genes which were highly expressed both in GRO and FGO (HS group) were highly enriched with H3K4me3 compared to genes in the LS, UP and DN groups (Figure S6C).

When we compared enrichments of H3K27me3 and H3K4me3 histone marks at promoters of affected genes to all genes with UP expression dynamics *in vivo*, we observed higher and lower enrichments, respectively, for all clusters regulated by CGI promoters in NGOs (Figures 6C; S6C) (Hanna *et al*., 2018b) as well as in E13.5 PGCs (Kawabata et al., 2019) and d6 PGCLCs (Kurimoto et al., 2015) (Figure S6D). These observations are consistent with our measurements of enrichment of CGI promoters with histone marks (Figure 6A) and suggested that CGI genes that are affected by the *in vitro* culture are enriched for PRC targets. Further, when we analyzed H3K27me3 enrichments in FGO and compared them to NGO oocytes (Figure 6C), we observed that CGI genes in clusters UP1-UP5 exhibited cluster specific changes in H3K27me3 enrichments, with genes with CGI promoters in UP2 showing a most noticeable removal of H3K27me3 from their promoters. In addition, CGI and non-CGI affected genes in DN group (i.e. DN1-DN5 clusters) showed various dynamics of H3K27me3 enrichment from NGO to FGO (Figure 6D). These observations suggest likely aberrant effects of PRC-mediated repression of genes belonging to UP2 and partially to UP3 clusters during the IVD treatment. We speculate, that genes belonging to UP2 and UP3 clusters were enriched for PRC1/2 targets in PGCs, PGCLCs as well as in NGOs and were subject to removal of repressive H3K27me3 histone marks to become up-regulated in the FGO *in vivo* (Figures 6E and S6I, J). The IVD treatment possibly caused instability and premature removal of these repressive marks from CGI promoters in UP group leading to premature activation of these genes.

Following this hypothesis, we investigated whether expression of components of PRC1 and PRC2 complexes was affected by *in vitro* oocyte development (Fig. 6F). Remarkably, we observed that the expression of several components of the canonical PRC1 complex was highly variable between growth conditions. For example, *Bmi1*/*Pcgf4* was downregulated in PGCLC versus PGC development while regulation of *Pcgf2* was labile in the IVD and IVG. The five *Cbx2*, *4*, *6*, *7*, and *8* genes were variably regulated under the *in vitro* culture as well. Likewise, components of variant PRC1 complexes such as *Yaf2*, *Kdm2b*, *Bcorl1*, *Skp2* and *Auts2* were to some extent also variably expressed. The role of such PRC1 complex members in the control of specific genes remains to be explored. Genetic loss-of-function studies revealed functional redundancy as well as critical roles of major PRC1 core components in PGC and oocyte development (Posfai *et al*., 2012; Yokobayashi et al., 2013). For members of the PRC2 complex, we observed deregulated expression of *Jarid2* and *Aebp2*, encoding two proteins involved in reading PRC1-catalyzed H2AK119ub1 and promoting PRC2 catalytic activity towards H3K27me3 (Kasinath et al., 2021). Expression of the *Kdm6b* H3K27me3 demethylase was also variable between culture conditions. Finally, *Bend3* encoding a protein implicated in PRC2 recruitment to paternal constitutive heterochromatin in early mouse embryos was significantly downregulated in the IVD and upregulated in the IVG (Saksouk et al., 2014). Taken together, we hypothesize that alterations in the expression of particular PRC2 and PRC1 components may impact the temporal dynamics of repression of target genes during *in vitro* germ cell development.

## Discussion

In this study, we recapitulate the differentiation of mouse PSCs to MII oocytes in culture with a comparable efficiency as previously reported (Hikabe *et al*., 2016). While the procedure yields many morphologically well-developed oocytes, competence to support early embryonic development is limited. Our analysis identifies the inability of parthenogenic eggs to initiate transcription as a major roadblock for successful embryonic development. Given the importance of nuclear PDH in regulating ZGA (Nagaraj *et al*., 2017), the reduced nuclear localization of PDH in PA embryos may also contribute to their low embryonic competence. Moreover, we observed abnormal acquisition of 5hmC and a failure of the STELLA protein to localize in the nucleus in BVSC-iPSC-derived parthenotes (Figure S4A, B). These data are compatible with a report that *Stella*-null embryos showed ectopic appearance of 5hmC in maternal chromatin, which induced abnormal accumulation of γH2AX and subsequent growth retardation (Nakatani et al., 2015). Ectopic acquisition of 5hmC in PSC-derived embryos likely results from TET3-mediated conversion of 5mC, even though 5mC levels in PSC-derived 2-cell embryos was comparable to that in *in vivo*-derived embryos (Figure 4F). Such differential response may in part be due to different sensitivity of antibodies for the respective epitopes. Possibly, reduced nuclear STELLA levels may impact on passive DNA demethylation. STELLA-mediated demethylation is achieved via inhibition of UHRF1 chromatin binding, thereby preventing DNMT1-mediated maintenance methylation, and is attenuated by nuclear export of STELLA (Du et al., 2019; Li et al., 2018). In mouse zygotes, cytoplasmic-localized STELLA undergoes ubiquitin-induced proteolytic cleavage (Shin et al., 2017). Proteosome-mediated degradation is incomplete and results in the association of a N-terminal STELLA fragment with early and re-cycling endosomal vesicles. Genetic experiments indicate that such cytoplasmic function is important for endo/exocytosis and required for pre-implantation development. Based on the reduced STELLA protein levels in iPSC derived zygotes, we speculate that factors and processes involving the ubiquitin-proteosome system or endo/exocytosis are in part deregulated upon *in vitro* oocyte generation. Since most *in vitro*-derived oocytes are morphologically indistinguishable from *in vivo*-derived oocytes, identification of intracellular abnormalities in *in vitro*-derived oocytes provides molecular markers for further improving the culture system.

### IVD culture step has a crucial effect on oocyte quality

Our data indicate that the low competence for preimplantation development results predominantly from IVD, and to lesser extent from IVG. For IVD, E12.5 gonads are dissociated, and gonadal somatic cells are mixed with d6 PGCLCs to form rOvaries. Transcriptional analyses have shown that d6 PGCLCs are comparable to migrating PGCs at E9.5 (Hayashi *et al*., 2012; Hayashi *et al*., 2011). Therefore, rOvaries consisting of d6 PGCLCs and E12.5 gonadal somatic cells contain developmentally heterochronic cell populations. Such heterochrony might cause aberrant follicle development during IVD. Nevertheless, our experimental data revealed no significant difference in developmental competence between iPSC-derived oocytes, developed in rOvaries consisting of d6 PGCLCs and E12.5 gonadal somatic cells, and E12.5 gonad-derived oocytes, developed in rOvaries consisting of E12.5 PGCs and E12.5 gonadal somatic cells (Figure 3; Table 1). Considering these data, not the heterochrony between d6 PGCLC and E12.5 gonadal somatic cells but rather the act of disrupting and reconstituting cellular interactions between PGCLCs and gonadal cells and culture conditions may impact on developmental competence.

PGCs develop into oocytes through formation of cysts and subsequently into follicles by close interaction with surrounding gonadal cells (Niu and Spradling, 2022; O’Connell and Pepling, 2021). In the IVD system, E12.5 gonads are dissociated, and gonadal somatic cells are mixed with PGCLCs to form rOvaries at day 8 of the culture. It is possible that components in gonads, especially the basement membrane and extracellular matrix (ECM), which contribute to achieve proper cell-cell interaction, were damaged or lost during this procedure. Transcriptome analysis using transplanted gonads/ovaries suggested development of GROs during the perinatal period are markedly subject to the ECM, which is also involved in oocyte dormancy (Nagamatsu *et al*., 2019). Our single oocyte transcriptome analysis revealed upregulation of genes involved in cell adhesion in GROs due to IVP and IVD and downregulation during IVG, possibly suggesting stage specific adaptation towards an altered extracellular environment (Figure S5E). Targeting the lack of extracellular components in rOvaries might improve the culture system and lead to *in vitro*-derived oocytes with higher embryonic competence after fertilization.

### Transcriptome comparison and modeling identifies differential expression of Polycomb target genes during IVD between *in vitro* and *in vivo* grown germ cells

In our study, we generated a large number of single oocyte RNA-Seq datasets as a resource for understanding the influence of each culture step on oocyte quality. For this, we benchmark the performance of *in vitro* and *in vivo* grown germ cells at different developmental stages. Our comparative expression analysis demonstrated that genes normally upregulated during oocyte growth are particularly vulnerable to *in vitro* culture conditions, leading to either aberrant up or down regulation in a development specific manner. Our epigenomic analysis indicates that many of such deregulated genes are normally controlled by PRC2 and PRC1 complexes catalyzing repressive H3K27me3 and H2AK119ub1 histone modifications thereby formatting repressive chromatin states. Since such modifications can also be removed by respective histone demethylases and de-ubiquitinating enzymes expressed in PGCs and oocytes (Figure 6F), *in vitro* culturing conditions likely result in altered expression of Polycomb target genes during oocyte growth. Genetic loss-of-function studies combined with spindle transfer experiments have identified a critical role for major PRC1 core components in oocyte development and specifying the maternal contribution required for zygotic transcription, timing of embryonic replication and embryonic development (Posfai *et al*., 2012). Moreover, Hikabe and colleagues reported that embryos obtained from *in vitro* generated oocytes were characterized by enlarged placentae (Hikabe *et al*., 2016). This finding is reminiscent of a failure in establishing non-canonical imprinted repression through PRC2-mediated H3K27me3 within the maternal genome in *in vitro* derived oocytes (Inoue *et al*., 2017a; Matoba et al., 2022). Hence, deregulation of Polycomb repression during *in vitro* oogenesis may directly or indirectly alter the maternal load of transcripts and proteins, as well as formatting the chromatin landscape in oocytes that normally confer embryonic competence, and possibly regulate ZGA upon fertilization.

The *in vitro* culture system for generating oocytes from PSCs has enormous potential for understanding germline development. Recently, follicles have been generated entirely from mouse PSCs without the use of donor tissues *in vitro* (Yoshino *et al*., 2021). Thus, wider application for mechanistic studies as well as new avenues in assisted reproduction are anticipated (Cyranoski et al., 2023; Saitou and Hayashi, 2021). Similar technology is already considered for obtaining human germ cells (Hwang et al., 2020; Irie et al., 2015; Sasaki *et al*., 2015; Yamashiro et al., 2018), and PGCLCs from endangered species to rescue animals from extinction (Hayashi et al., 2022). Our work provides molecular insights into *in vitro* oogenesis and identifies critical steps to direct efforts for future improvement.

## Experimental Procedures

### Mice

C57BL/6J, CAST/EiJ and DBA/2J mice were purchased from Jackson Laboratory. Swiss Webster and 129S6/SvEvTac mice were purchased from Taconic Biosciences. C57BL/6J females were mated with DBA/2J males to obtain hybrid mice (B6D2F1). All mice were housed in the animal facility of ETH Zurich. All animal experiments were performed under the license ZH152/17 in accordance with the standards and regulations of the Cantonal Ethics Commission Zurich.

### Cell lines

C57BL/6J females were mated with 129S6/SvEvTac males to obtain hybrid embryos to establish ESC lines. After genotyping of sex (Aizawa et al., 2020), a female ESC line was co-transfected with a piggyBac vector carrying a CAG-EGFP-IRES-hygro transgene and a hyperactive piggyBac transposase expression plasmid (Yusa et al., 2011) using lipofectamine 2000 by following a manufacturer’s protocol. Subsequently, single cells with EGFP expression were sorted by FACS. After cell growth, a single colony was used to establish the GFP-ESC line. The female BVSC-ESC line bearing Blimp1-Venus and Stella-ECFP transgenes was established as previously described (Hayashi *et al*., 2012). The female BVSC-iPSC line established from adult tail tip fibroblasts, also called iPS TTF_4FC6 (Hikabe *et al*., 2016), was a gift from Katsuhiko Hayashi. All these 3 cell lines were maintained feeder-free on ornithine- and laminin-coated plates using 2i+LIF medium (Hayashi and Saitou, 2013).

### *In vitro* PGC differentiation (IVP)

Differentiation of PSCs into EpiLCs and PGCLCs was induced by following a previously published protocol with a few modifications (Hayashi and Saitou, 2013). For EpiLC differentiation, 3.4 x 10^5^ PSCs per well were plated on a 6-well plate coated with 16.7 mg/ml human plasma fibronectin in EpiLC medium. For PGCLC differentiation, 2.25 x 10^5^ EpiLCs per well were plated in a Spherical plates 5D plate (Kugelmeiers) with PGCLC medium containing BMP4 (500 ng/ml), SCF (100 ng/ml), LIF (1,000 IU/ml) and EGF (50 ng/ml) without BMP8a. The medium was changed at day 1 with EpiLC medium, and at day 6 with GK15 medium supplemented with SCF (50 ng/ml), LIF (500 IU/ml) and EGF (25 ng/ml). Images of cells were acquired using an Olympus MVX10 Stereo-Zoom microscope equipped with an Olympus DP73 camera using the cellSens software.

### *In vitro* oocyte differentiation (IVD)

The IVD culture condition was adapted from previously published protocols with some modifications (Hayashi et al., 2017; Hikabe *et al*., 2016). At day 8 of the culture, PGCLCs, positive for both Blimp1-Venus and Stella-ECFP or for both SSEA1 and integrin β3, were sorted using a FACSAria III (BD Bioscience). The collected PGCLCs were resorted once more for purification using the same gate by FACSAria III. At the same time, E12.5 female embryonic gonads, derived from outbred Swiss Webster mice, were harvested. To isolate gonadal somatic cells, PGCs were depleted by magnetic-activated cell sorting using both SSEA1 and CD31 antibodies coupled to magnetic beads (130-094-530 and 130-097-418, Miltenyi Biotech) in accordance with the manufacturer’s protocol. PGCLCs were aggregated with isolated gonadal somatic cells in a low-binding 96-well plate (174929, Thermo Scientific) for 2 days in GK15 medium supplemented with 1 μM retinoic acid. 5,000 PGCLCs and 50,000 gonadal somatic cells were cultured to produce one rOvary. At day 10 of the culture, rOvaries were placed on Transwell-COL membranes (3492, Corning) in a 6-well plate, in which the membrane is contacted with the surface of α-IVD medium composed of αMEM supplemented with 2% FBS (A3161001, Thermo Fisher), 150 µM ascorbic acid (Merck), 55 µM β-mercaptoethanol (Thermo Fisher), 2 mM glutamax (Thermo Fisher) and penicillin-streptomycin (Thermo Fisher). At day 12 of the culture, rOvaries were soaked in the medium by adding 2 ml of α-IVD medium per a well. At day 14, half of medium was changed to Stem-IVD medium composed of StemPro-34 SFM (Thermo Fisher) supplemented with 10% FBS, 150 µM ascorbic acid, 55 µM β-mercaptoethanol, 2 mM glutamax and penicillin-streptomycin. At day 17, the medium was replaced with Stem-IVD medium supplemented with 600 nM fulvestrant (Merck). At day 21, the medium was changed to Stem-IVD medium without fulvestrant. Half of the medium was changed at day 12, 19, 21, 23, 25, 27 and 29.

For comparing the development, E12.5 gonad-derived rOvaries were also prepared. We dissociated E12.5 gonads and then aggregated all gonadal cells including PGCs in low-binding plates to reconstitute rOvaries for 2 days in GK15 medium supplemented with 1 μM retinoic acid. 55,000 gonadal cells were used to produce one rOvary. Then, rOvaries were placed on Transwell-COL membranes by following the similar IVD protocol as PSC-derived rOvaries for 21 days.

Samples were imaged under the microscope (Axio Observer Z1, Zeiss) equipped with an ORCA-Flash4.0 camera (Hamamatsu Photonics K.K.). Images were processed using Zeiss Zen Pro 2.0 software.

### *In vitro* growth (IVG)

The IVG culture condition was adapted from previously published protocols with a few modifications (Hayashi *et al*., 2017; Hikabe *et al*., 2016). At day 31 of the culture, follicles in rOvaries were mechanically dissociated using 30G needles. Dissociated follicles were kept on the Transwell-COL membranes contacted with IVG medium composed of αMEM supplemented with 2% polyvinylpyrrolidone (PVP360, Merck), 5% FBS, 100 mIU/ml follicle-stimulating hormone (FSH) (Puregon, MSD), 150 µM ascorbic acid, 55 µM β-mercaptoethanol, 2 mM glutamax, 55 µg/ml sodium pyruvate and penicillin-streptomycin. At day 33, follicles were incubated in 0.1% collagenase type I (Worthington Biochemicals) for 15 minutes. After washing with αMEM supplemented with 5% FBS 3 times, the transwell was placed into a new 6-well plate filled by IVG medium with the same height of the membrane. From day 31 to 34, the IVG medium was supplemented with 15 ng/ml BMP15 (ab127067, abcam) and 15 ng/ml GDF9 (739-G9-010, R&D Systems). At day 34, follicles were soaked in the medium by adding 2 ml of IVG medium per well. Half of the medium was changed at day 36, 38, 40 and 42.

Follicles dissected from E12.5 gonad-derived rOvaries and from 6 dpp ovaries were also subjected to the IVG at day 23 of the culture and after the dissection of 6 dpp ovaries, respectively. The same IVG protocol including the collagenase treatment was applied to 4-10 follicles dissected from the rOvaries or from 6 dpp ovaries as follicles from PSC-derived rOvaries. The duration of the IVG was 11-12 days, which were 1-2 days shorter than that for PSC-derived follicles since dying cells emerged in several follicles when they were cultured for 13 days.

### *In vitro* maturation (IVM)

The IVM culture condition was adapted from previously published protocols (Hayashi *et al*., 2017; Hikabe *et al*., 2016). At day 44 of the culture, oocytes with surrounding granulosa and cumulus cells were harvested from expanded follicles with a diameter of roughly over 200 µm at the longest axis using a fine glass capillary. These complexes were transferred to IVM medium composed of αMEM supplemented with 5% FBS, 100 mIU/ml FSH, 4 ng/ml EGF (315-09, Peprotech), 1.2 IU/ml human chorionic gonadotropin (HCG) (C1063, Merck), 25 µg/ml sodium pyruvate and penicillin-streptomycin. At 16 hours of the culture, swollen cumulus-oocyte complexes (COCs) were subjected to IVF or parthenogenetic activation. The same IVM protocol was applied to expanded follicles derived from E12.5 gonads and 6 dpp ovaries at day 34-35 and at day 11-12 of the culture, respectively.

### *In vitro* fertilization (IVF) and preimplantation development

Spermatozoa were collected from the cauda epididymis of male mice. B6D2F1 males were used for assessment of preimplantation development, and CAST/EiJ males were used for transcription analysis of embryos. Collected spermatozoa were capacitated by incubation for 1 hour in Sequential Fert (83020010, ORIGIO). After capacitation, spermatozoa were incubated with COCs after the IVM in Sequential Fert for 6 hours. The zygotes were collected and transferred to KSOM medium (MR-020P, Merck) for preimplantation development. After 2 days of the culture, embryos were transferred to fresh KSOM medium. The embryos were counted every single day to measure their developmental ratio. Statistical analysis was performed by GraphPad Prism 8 software using an unpaired t test.

### Parthenogenetic activation

COCs were placed in M2 medium (M7167, Merck) and stripped from cumulus cells by treating with 0.1% hyaluronidase (H4272, Merck). MII oocytes were identified by their morphology with first polar body extrusion. All the oocytes harvested from COCs were transferred to activation medium composed of KSOM medium supplemented with 5 mM strontium chloride (13909, Merck) and 2 mM EGTA (A0878, AppliChem) (Kishigami and Wakayama, 2007). After 6 hours, activated embryos were transferred to KSOM medium for subsequent preimplantation development. After 2 days of the culture, embryos were transferred to fresh KSOM medium.

### Assessment of follicle development

After the IVG culture, expansion of follicles derived from BVSC-iPSC, E12.5 gonad and 6 dpp follicle was assessed at day 44, day 34 and day 12 of the culture, respectively. Since follicles were not round, the longest part in each follicle was measured as a diameter under a stereomicroscope (SMZ745, Nikon) with an eyepiece reticle. Each follicle was categorized into one of 3 groups depending on its diameter: 0 - 200 µm; 200 - 400 µm; over 400 µm.

### Genotyping

DNA extraction of cells was performed as previously described (Aizawa *et al*., 2020). PCR was performed using Phusion Hot Start II DNA Polymerase (Thermo Fisher Scientific) following the manufacturer’s protocol. PCR products were separated by electrophoresis on 1.5% agarose gels and stained with ethidium bromide for visualization under a UV transilluminator. The following primer sequences were used for genotyping of sex and cell line identification: Blimp1-mVenus-5, 5’-ACT CAT CTC AGA AGA GGA TCT G-3’; Blimp1-mVenus-3, 5’-CAC AGT CGA GGC TGA TCT CG-3’; Prdm14 WT-5, 5’-AAG GTT CTG GGA ACT GGA TGT C-3’; Prdm14 WT-3, 5’-CAC AAT ATG CTG GCA TGC GTT C-3’; Stella-ECFP-5, 5’-CGA GCT AGC TTT TGA GGC TT-3’; Stella-ECFP-3, 5’-AAC TTG TGG CCG TTT ACG TC-3’; SRY2, 5’-TCT TAA ACT CTG AAG AAG AGA C-3’; SRY4, 5’-GTC TTG CCT GTA TGT GAT GG-3’; Xist-14, 5’-GTA GAT ATG GCT GTT GTC AC-3’; Xist-16, 5’-CTC CAT CCA AGT TCT TTC TG-3’.

### Immunostaining analysis

PSC-derived 2-cell embryos were prepared by parthenogenetic activation of oocytes after IVM. Control 2-cell embryos were prepared by activation of oocytes harvested from superovulated C57BL/6J females. Control 1-cell zygotes were prepared by IVF of oocytes and spermatozoa harvested from superovulated C57BL/6J females and B6D2F1 males, respectively.

For immunostaining of histone marks and STELLA, 2-cell embryos were collected at 48 hours after HCG injection or the start of IVM. 2-cell embryos for immunostaining of PDH were collected at 44 hours after HCG injection or the start of IVM. Immunostaining was performed by following a published protocol (Nagaraj *et al*., 2017). Embryos were fixed in 4% paraformaldehyde in PBS for 30 min at room temperature, permeabilized for 30 min in PBS with 0.4% Triton X-100 (PBST4), blocked in PBST4 with 3% bovine serum albumin (PBST4A) for 30 minutes and incubated with the desired primary antibody in PBST4A overnight at 4C. The embryos were washed in PBST4 four times for 10 min each, blocked with PBST4A, and incubated with the appropriate secondary antibody and DAPI overnight at 4C. The embryos were washed 3 times for 10 min each in PBST4, then deposited on glass slides and mounted in Vecta-shield (Vector Laboratories).

BrUTP incorporation assay was performed by following a published protocol (Suzuki et al., 2015) with a few modifications. 2-cell embryos were collected at 53 hours after HCG injection or the start of IVM. BrUTP incorporation was performed by electroporation using the Super Electroporator NEPA 21 (NEPAGENE) as previously described (Dumeau et al., 2019). Embryos were washed in PBS and then transferred in a line on the glass chamber between electrodes filled with PBS containing 10 mM BrUTP (Merck). The poring pulse (voltage: 30 V, pulse length: 3 ms, pulse interval: 100 ms, number of pulses: 6, +) and the transfer pulse (voltage: 5 V, pulse length: 50 ms, pulse interval: 50 ms, number of pulses: 5, ±) were applied. The embryos were washed twice and cultured in KSOM for 1 hour. Subsequently, fixation and immunostaining of embryos followed the protocol for histone marks or PDH described above. Anti-BrdU antibody (B8434, Merck) was used as the primary antibody.

For immunostaining of 5mC and 5hmC, 2-cell embryos were collected at 48 hours after HCG injection or start of IVM. Immunostaining was performed by following a published protocol (Nakamura *et al*., 2007) with a few modifications. Embryos were treated with 2 M HCl for 20 min and subsequently washed with 0.05% Tween 20 in PBS (PBST5) after permeabilization. The embryos were blocked for 1 h in 1% bovine serum albumin and 0.05% Tween 20 in PBS (PBST5A), and then incubated overnight in anti-5mC and anti-5hmC antibodies (NA81-50UG, Millipore; 39069, Active Motif). The following day the embryos were washed in PBST5 four times for 10 min each, blocked with PBST5A, and incubated with the appropriate secondary antibody and DAPI overnight at 4C. Embryos were washed 3 times for 10 min each in PBST5, then deposited on glass slides and mounted in Vecta-shield.

After immunostaining of embryos, images were captured using a Leica TCS SP8 confocal microscope equipped with a sCMOS camera (Hamamatsu Orca Flash 4.0). Processing and quantification of images were performed using Fiji software (https://imagej.net/software/fiji/). For quantification of fluorescent signals, regions of nuclei and cytoplasm of 2-cell embryos were manually selected using DAPI signals and bright field images. The mean intensity of two nuclei was subtracted with the mean intensity of cytoplasm of two blastomeres in each 2-cell embryo. Quantified data was compiled and analyzed by GraphPad Prism 8 software using an unpaired t test. Data was considered significant if p < 0.05.

### Isolation and sequencing of PGC and PGCLC transcriptomes

E12.5 PGCs and BVSC-iPSC-derived PGCLCs were used for their transcriptome analysis with bulk RNA-seq (Figure 5A). To prepare pooled PGCs, E12.5 female embryonic gonads were harvested from inbred C57BL/6J fetuses. PGCs, positive for both SSEA1 and integrin β3, were collected using a FACSAria III (BD Bioscience). D6 PGCLCs derived from a BVSC-iPSC line were also collected at day 8 of the culture by following the same protocol as the start of the IVD culture. Four replicates of respective PGC and PGCLC pools, consisting of 30,000 to 340,000 cells, were prepared from independent experiments. Total RNA was extracted using RNeasy Mini Kit (Qiagen) according to the manufacturer’s protocol, including removal of genomic DNA. The RNA quality was determined by 2200 TapeStation (Agilent Technologies). All samples had a RIN value of greater than 8. Extracted RNA was prepared for sequencing using the TruSeq Stranded mRNA (Illumina) following the manufacturer’s protocol. Briefly, 100 to 1000 ng of total RNA was poly-A-enriched and reverse-transcribed into double-stranded cDNA. The cDNA samples were fragmented, end-repaired, and adenylated before ligation of TruSeq adapters. Fragments containing TruSeq adapters for multiplexing on both ends were selectively enriched with PCR. Sequencing was performed on an Illumina NovaSeq 6000 (Illumina), with sequencing depth of 20 million reads per sample and sequencing configuration of single-end 100 bp.

### Isolation and sequencing of single oocyte transcriptomes

RNA sequencing libraries for single oocytes were prepared according to the Smartseq2 protocol (Picelli et al., 2014). For each experimental condition, 20-25 independent libraries were prepared. Firstly, each GRO and FGO derived from BVSC-iPSCs, E12.5 gonads, 6 dpp follicles and *in vivo* samples (Figure 5A) was dissociated from follicular somatic cells using 30G needles, followed by treatment with Accumax (Innovative Cell Technologies) for 5 min and by pipetting to remove somatic cells completely. The zona pellucida of each GRO and FGO was removed by acid tyrode’s solution (Merck) and samples were individually washed in PBS supplemented with 0.01% PVA (PBS-PVA) and lysed in the lysis buffer composed of 0.09% Triton-X 100, 2 U SUPERase IN RNase inhibitor (Invitrogen, AM2694), 2.5 μM Oligo-dT primer (Microsynth AG), dNTP mix (2.5 mM each, Promega), ERCC RNA Spike-In Mix (1 : 3.2 x 10^7^, Thermo Fischer Scientific 4456740) in individual tubes of a 8-well strip, then immediately frozen on the dry ice and kept at -80C for longer storage. Next, sample lysate was denatured at 72C for 3 min and quickly chilled on ice. The reverse transcription mix was composed of 100 U SuperScript II reverse transcriptase (Thermo Fischer Scientific, 18064014), 5 U SUPERase IN RNase inhibitor (Thermo Fischer Scientific, AM2696), 1X Superscript II first-strand buffer, 5 mM DTT (provided with SuperScript II reverse transcriptase), 1 M Betaine (Sigma, B0300-1VL), 6 mM MgCl2, 1 μM template-switching oligos (TSOs) (“AAGCAGTGGTATCAACGCAGAGTACATrGrG+G”; Exiqon) was added to obtain a total volume 10 μl and the reverse transcription was performed in PCR machine. Then PCR pre-amplification was performed in a total volume of 25 μl by adding 1X KAPA HiFi HotStart Ready Mix (KAPA Biosystems, KK2602), 0.1 μM ISPCR primers (Microsynth AG). The preamplification PCR cycle numbers were 15-16. Pre-amplified cDNA was purified with SPRI AMPure XP beads (Beckman, sample to beads ratio 1:1) and eluted in 15 μl Buffer EB (QIAGEN). 1 ng of pre-amplified cDNA was used for the tagmentation reaction (55C, 7 min) using a homemade Tn5 tagmentation mix (1x TAPS-DMF buffer, Tn5-transposase) in total volume of 20 μl. The reaction was stopped by adding 5 μl of 0.2% SDS (Invitrogen, 24730020) and kept at 25C for 7 min. Adapter-ligated fragment amplification was done using Nextera XT index kit (Illumina) in a total volume of 50 μl (1x Phusion HF Buffer, 2 U of Phusion High Fidelity DNA Polymerase (Thermo Fischer Scientific, F530L), dNTP mix (0.3 mM each, Promega) with 9-10 cycles of PCR. Library was purified by SPRI AMPure XP beads (sample to beads ratio 1:1) and eluted in 12 μl Buffer EB. Sequencing was performed on an Illumina HiSeq 2500 machine with single-end 75-bp read length (Illumina).

### Transcriptional analysis

#### Alignment and of RNA-Seq data

RNA-Seq datasets were aligned to a custom genome containing the *Mus musculus* genome assembly (GRCm38/mm10 Dec. 2011) and ERCC92 sequences using STAR (Dobin et al., 2013) with parameters “-outFilterMultimapNmax 300 -outMultimapperOrder Random -outSAMmultNmax 1 -alignIntronMin 20 -alignIntronMax 1000000”, allowing multimappers with up to 300 matches in the genome and choosing positions for multimappers randomly (Table S5).

#### Selection of exonic and promoter regions for genes and quantification of gene expression

A random transcript isoform for each gene was chosen from Bioconductor annotation package TxDb.Mmusculus.UCSC.mm10.knownGene (version 3.2.2) and promoter regions were constructed as regions 1.5 kb upstream and downstream from transcription start site. Any overlaps between promoters were removed by choosing a random promoter among promoters which overlap. Resulting sets of non-overlapping promoter and exonic genomic regions for the random transcript isoforms for each gene were used in further analyses.

Expression quantification for selected exonic genomic regions was done using QuasR R package (Gaidatzis et al., 2015) selecting only uniquely mapped reads (mapqMin=255). RPKM values for genes were calculated by normalizing exonic read counts to total exonic length of each gene and total number of reads mapping to all exonic regions in each library. RPKM values were log2 transformed using formula log2(RPKM + psc) – log2(psc) where pseudo-count psc was set to 0.1.

#### Principal component analysis of PGCLC and PGC samples

To compare RNA-Seq data for day 6 PGCLC and E12.5 female PGCs generated for this study with previously published datasets we downloaded from the GEO repository RNA-seq datasets for E12.5 female PGC (GSE87644) (Ohta *et al*., 2017) and for day 6 PGCLC (GSE67259) (Sasaki *et al*., 2015). Relative to mean expression values for each study separately were calculated by subtracting average log2(RPKM) values across samples. Using relative expression values for samples generated for this study and for previously published datasets the Principal Component Analysis (PCA) was done using R function *prcomp* (Figure S5A).

#### Differential expression analysis of PGCLC and PGC samples

R package *edgeR* (McCarthy et al., 2012) was used to study gene expression differences between day 6 PGCLC and E12.5 female PGCs both for previously published and generated for this study datasets. Generalized Linear Model was fit using cell type (PGC or PGCLC) as covariates. Statistical significance was estimated using log-likelihood tests and the Benjamini-Hochberg method was used to correct for multiple testing.

#### Estimating of gene expression responses to stages of in vitro development

To minimize potential biases from outlier samples we selected samples based on oocyte sizes for further analysis (Table S5). Genes with CPM higher than 1 in at least 3 samples were used for the analysis. Samples for GROs and FGOs were separated and for each stage the gene expression responses were estimated by fitting generalized linear model with edgeR using model matrix encoding stages of *in vitro* or *in vivo* development underwent by each sample, i.e. IVP for oocytes in cohorts A2, A3 or *in_vivo* for oocytes in cohorts B2, B3, C3, D2 and D3; IVD for oocytes in cohorts A2, A3, B2, B3 or *in_vivo* for cohorts C3, D2 and D3; IVG for cohorts A3, B3, C3 and *in_vivo* for cohort D3. The expression responses which in this model are represented by coefficients for IVP, IVD and IVG for each gene were fit and statistical significance was estimated using log-likelihood test and corrected for multiple testing using the Benjamini-Hochberg method.

#### Gene ontology enrichment analysis

Enrichment analysis for Gene Ontology (GO) terms was done using R package *topGO* (version 2.48.0) (Alexa and Rahnenfuhrer, 2022) with parameters *method=”weight01”* and *statistic=”fisher”* extracting GO gene annotation from the Bioconductor Annotation Package *org.Mm.eg.db* (version 3.15.0) (Carlson, 2022).

Visualization of GO enrichments was done by calculation of pairwise Jaccard distances between significant GO terms based on intersections and unions of significantly affected gene sets having corresponding GO term annotations. After pairwise Jaccard distances between GO terms were calculated we applied multidimensional scaling (MDS) using R function *cmdscale* and represented GO terms on a 2D plot where size was scaled by obs./exp. ratio, color was chosen to reflect statistical significance and relative position reflects similarities in gene sets (Figures S5 D,E).

#### Investigation of promoter features of affected genes

Previously selected promoter regions were classified into CGI and non-CGI promoters by calculating observed versus expected ratios of CpG dinucleotides and fitting gaussian mixture model with 2 components.

Enrichment analysis for transcription factor motifs at promoter regions was performed using function *calcBinnedMotifEnrR* in R package *monaLisa* (Machlab *et al*., 2022) for which genes were binned according to combination of type of expression response (up-regulated or down-regulated) and type of expression dynamics *in vivo* (LS, UP, DN and HS groups in Figure 5C) for each stage of *in vitro* development separately (IVP@GRO, IVD@GRO, IVP@FGO, IVD@FGO and IVG@FGO). As background set we used option “otherBins” for *calcBinnedMotifEnrR* function. We used a set of position weight matrices (PWMs) from the *JASPAR2020* Bioconductor R package for vertebrates which encompasses 746 PWMs.

#### Alignment and analysis of ChIP-seq data

Previously published ChIP-Seq or Cut&Run samples for PGCLC as well as for different stages of *in vivo* development were downloaded from the GEO repository (see Table S5 for list of published datasets and accession IDs used in this study). The quality of the data was assessed using FastQC (v0.11.8) and adapters were trimmed using TrimGalore (v0.6.2) (Krueger, 2015). All datasets were aligned to mm10 mouse genome using STAR with parameters “-alignIntronMin 1 -alignIntronMax 1 –alignEndsType EndToEnd - alignMatesGapMax 1000 -outFilterMatchNminOverLread 0.85 –outFilterMultimapNmax 300 -outMultimapperOrder Random -outSAMmultNmax 1” and possible PCR duplicates were removed using samtools (Li et al., 2009).

Quantification for previously selected promoter regions were done using QuasR R package (Gaidatzis *et al*., 2015) selecting only uniquely mapped reads (mapqMin=255). Log2(CPM) values calculated by adding a pseudo-count of 16 to promoter read counts, normalizing to total number of mapped reads in each sample and log2 transformation.

## Acknowledgments

We are grateful for the advice on the *in vitro* culture protocol provided by Orie Hikabe and Katsuhiko Hayashi (Osaka University, Japan). A BVSC-iPSC line was kindly provided by Katsuhiko Hayashi. We thank the ETH Phenomics Center (ETH Zurich), Flow Cytometry Core Facility (ETH Zurich), Scientific Center for Optical and Electron Microscopy (ETH Zurich), and Functional Genomics Center Zurich for their technical assistance. We thank Sebastien Smallwood and colleagues (Functional Genomics Platform FMI, Basel) for technical assistance. This study was supported by an EMBO Short Term Fellowship (ASTF 207-2016) and Swiss National Science Foundation (grant 31003A_152814/1). This research has received funding from the Novartis Research Foundation and the European Research Council (ERC) under the European Union’s Horizon 2020 research and innovation programme (grant agreement ERC-AdG 695288 - Totipotency).

## Author Contributions

E.A., E.A.O., A.H.F.M.P., and A.W. conceptualized experiments. E.A., Y.K.K., and C.-E.D. performed data collection and/or analysis. E.A.O. performed bioinformatic analysis. E.A., E.A.O., A.H.F.M.P., and A.W. wrote the manuscript. S.N. and M.S. provided training, reagents and proofread the manuscript.

## Supplemental Information

Figures S1-S6 Table S1-S7

## Supplemental Table Legends

**Table S1.** Summary of non-follicle development in rOvaries

| Cell line | Cells sorted | All rOvary | rOvary with only follicle development<br>(% of all rOvary) | rOvary with non-follicle development<br>(% of all rOvary) |
| --- | --- | --- | --- | --- |
| GFP-ESC | SSEA1 + Integrin-β3 + | 42 | 0<br>(0%) | 42<br>(100%) |
| BVSC-ESC | SSEA1 + Integrin-β3 + | 7 | 0<br>(0%) | 7<br>(100%) |
| BVSC-ESC | Blimp1-Venus + Stella-CFP + | 24 | 20<br>(83.3%) | 4<br>(16.7%) |
| BVSC-iPSC | Blimp1-Venus + Stella-CFP + | 980 | 818<br>(83.5%) | 162<br>(16.5%) |

Table S2. Expression differences between in vitro d6 PGCLCs and E12.5 PGCs as determined by bulk RNA-seq experiment in this study and previously by RNA-seq experiments in (Ohta et al., 2017; Sasaki et al., 2015).

Table S3. GO term enrichment analysis for up-regulated and down-regulated genes between d6 PGCLCs and E12.5 PGCs as determined by bulk RNA-Seq experiment in this study.

Table S4. Read statistics and mapping rates for single oocyte RNA-seq samples (sheet “Single Oocyte RNA-Seq”), bulk RNA-Seq samples for day 6 PGCLC and E12.5 female PGC (sheet “Bulk PGCLC and PGCLC RNA-Seq”) and GEO/DDBJ accession IDs for published genomic datasets used in this study (sheet “Published genomics datasets”).

Table S5. Gene expression responses to modeled effects of each step of *in vitro* development as revealed by modeling of gene expression in A2, B2 D2 GRO cohorts and A3, B3, C3 and D3 FGO cohorts (sheet “Gene expression responses”) as well as gene expression aggregated across oocyte cohorts (sheet “Aggregated gene expression”).

Table S6. GO term enrichment for genes with significant response to in vitro development.

Table S7. Results of enrichment analysis of transcription factor motifs within promoters of genes with particular expression dynamics from GRO to FGO in vivo and affected by each stage of in vitro development. Columns Motif and Motif.ID represent motif names and ID in the JASPAR2020 database; Effect and geneGroup reflect names of gene groups on which the test was run; log2Enr, negLog10Pval and negLog10Pval.adj represent enrichments, - log10(p-values) and -log10(adjusted p-values); NumHits represents number of promoters with a corresponding motif and total number of promoters in a group.

**Figure S1.**
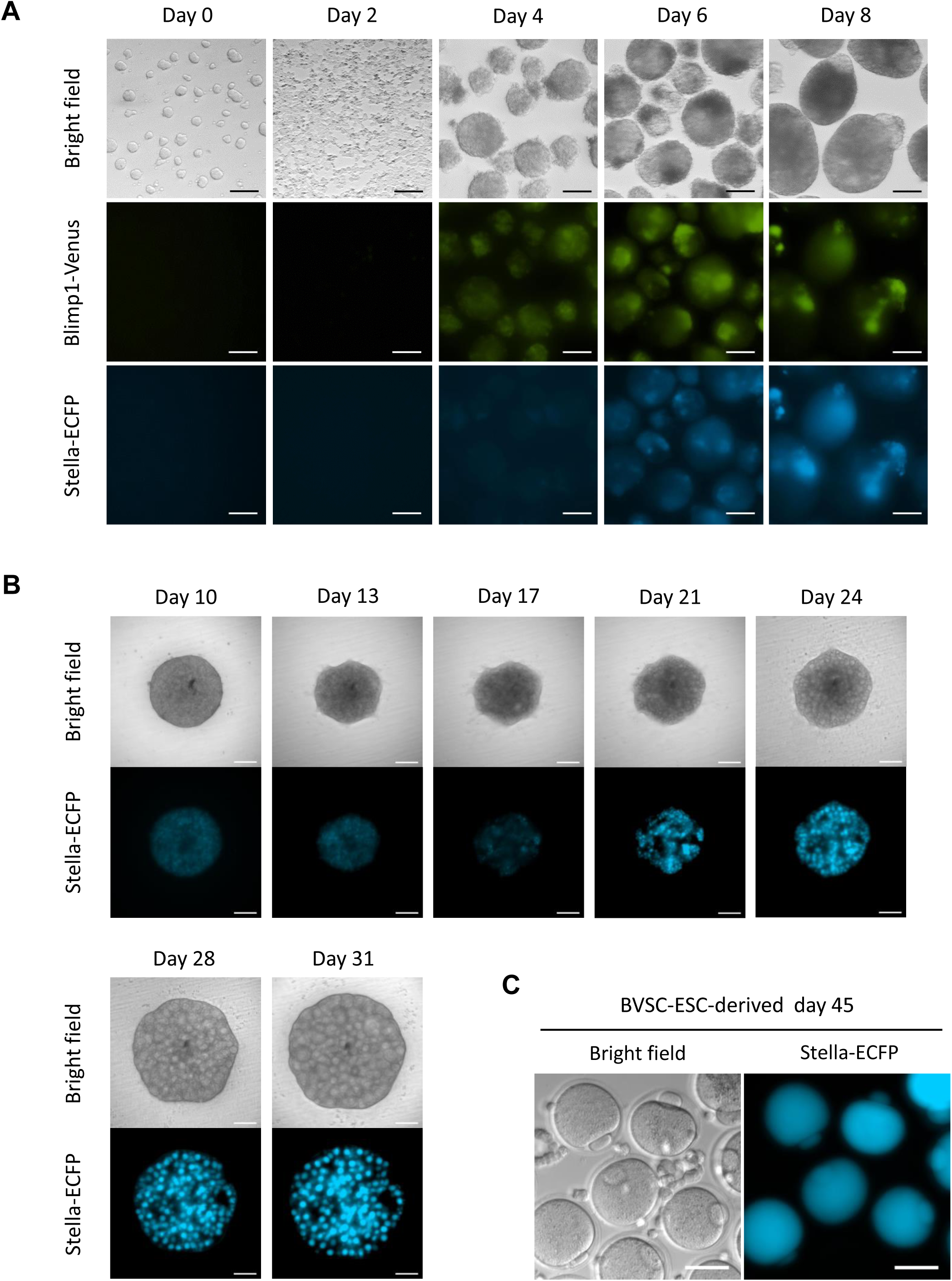
PGCLC differentiation and development of oocytes derived from BVSC-ESCs. (A) Differentiation of BVSC-ESCs to PGCLCs. Expression of Blimp1-Venus and Stella-ECFP as well as bright field images are shown from day 0 to 8 of the culture. Blimp1-Venus and Stella-ECFP started their expression at day 4 and 6, respectively. Bar scale, 100 µm. (B) Development of a BVSC-ESC-derived rOvary from day 10 to 31 of the culture. Stella-ECFP expression transiently decreased around day 17, followed by emergence of round oocytes expressing Stella-ECFP. Bar scale, 100 µm. (C) MII oocytes containing first polar bodies derived from BVSC-ESCs at day 45 of the culture. Oocytes without Stella-ECFP expression were presumably derived from PGCs in E12.5 gonads contaminated at day 8 of the culture. Bar scale, 100 µm.

**Figure S2.**
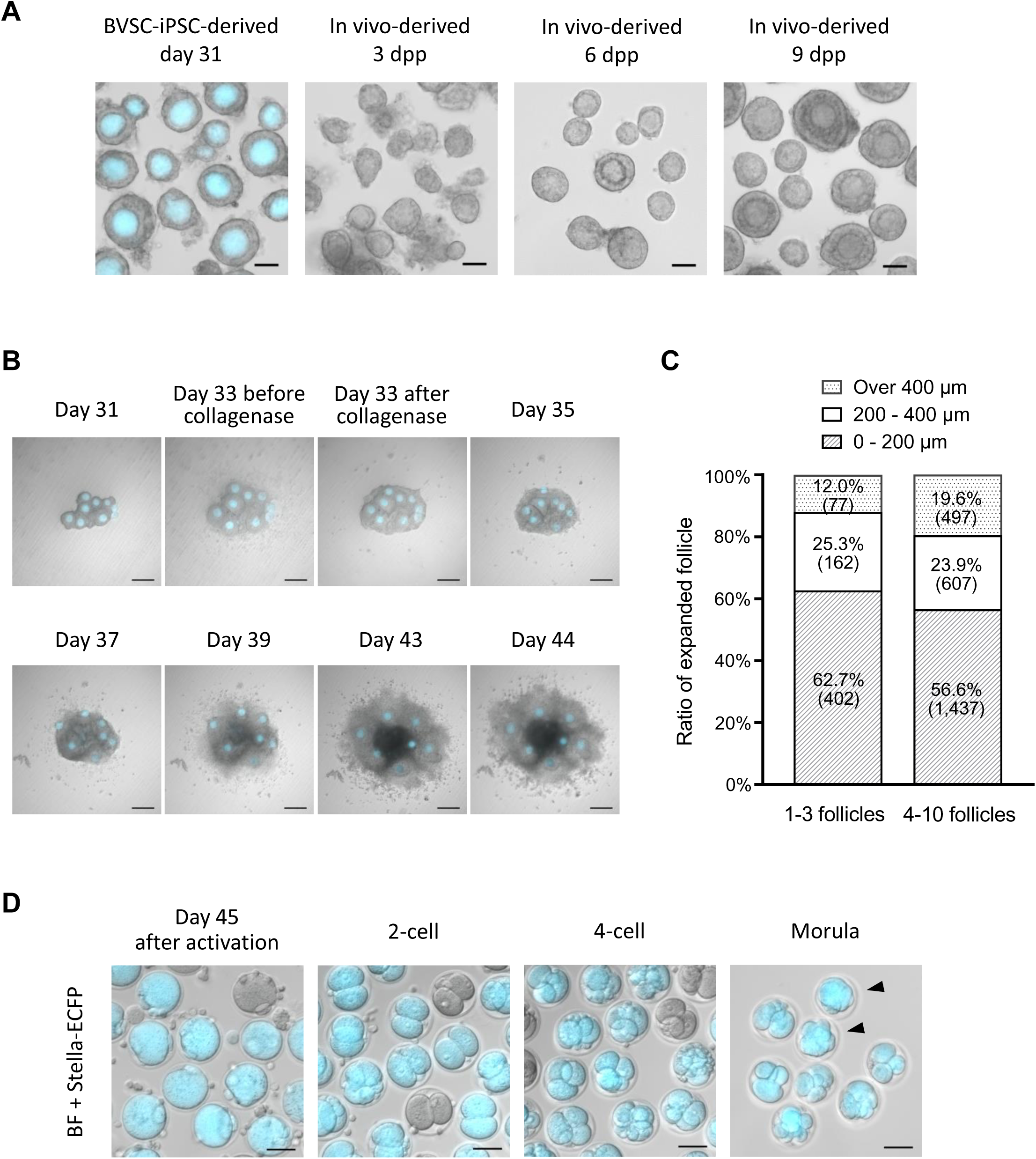
Development of BVSC-iPSC-derived follicles and preimplantation development of parthenotes. (A) A representative morphology of BVSC-iPSC-derived follicles at day 31, and *in vivo*-derived follicles at 3, 6 and 9 dpp. Stella-ECFP expression was merged with the bright field image of BVSC-iPSC-derived follicles. Bar scale, 50 µm. (B) IVG of BVSC-iPSC-derived follicles isolated from a rOvary. Bright field images were merged with Stella-ECFP expression. Cultured follicles were treated with collagenase at day 33 of the culture. Some follicles expansively developed at day 43. Scale bar, 200 µm. (C) Size of developed follicles at day 44 of the culture. The largest diameter of BVSC-iPSC-derived follicles in 2 conditions were measured; 1-3 follicles (left) and 4-10 follicles (right) placed at the same location during IVG. Brackets represent number of counted follicles. The data of 4-10 follicles is identical to the data of iPSC-derived follicle in Figure 3A. (D) Preimplantation development of BVSC-iPSC-derived oocytes after parthenogenetic activation. Embryos without Stella-ECFP expression presumably developed from E12.5 PGCs, which were possibly mixed with gonadal somatic cells for co-culture at day 8. Arrowheads indicate morulae. Bar scale, 50 µm.

**Figure S3.**
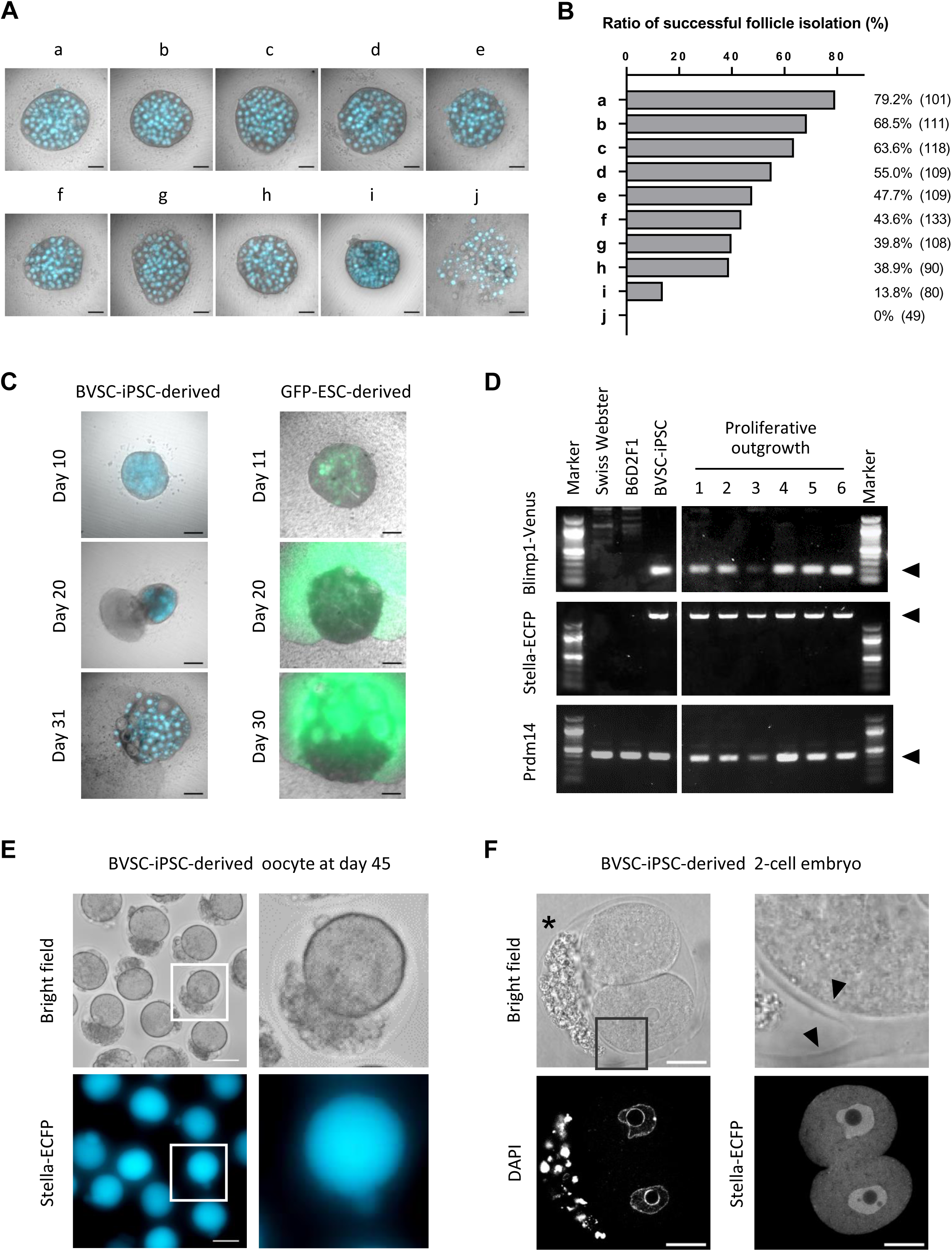
Assessment of FBS and abnormal development of rOvaries/oocytes. (A) Representative morphology of rOvaries at day 31 of the culture. Nine FBS and one serum replacement (a - j) were tested to assess follicle formation by the IVD culture. Bright field images merged with Stella-ECFP are shown. The commercial companies and catalog numbers of respective FBS and a serum replacement are as follows: a, Life Technologies, A3161001; b, Life Technologies, A3160801; c, GE Healthcare, SH30071.02; d, GE Healthcare, SV30160.02; e, Sigma, F0926; f, Life Technologies, A3160901; g, PAN Biotech, P30-1702; h, GE Healthcare, SH30084.02; i, Life Technologies, 10828-028; j, Equitech-Bio, SBSU30-0500. (B) At day 31 of the culture, each rOvary was mechanically dissected by 30G needles to isolate single secondary follicles. The ratio of successful follicle isolation was calculated based on the number of isolated single secondary follicles divided by the number of secondary follicles attempted to isolate, which are shown in brackets. The data is based on two independent experiments. (C) Development of proliferative outgrowth from rOvaries. Bright field images were merged with Stella-ECFP (left) and CAG-GFP (right). PGCLCs positive for both Blimp1-Venus and Stella-ECFP or for both SSEA1 and integrin-β3 were used for BVSC-iPSC-derived (left) or GFP-ESC-derived (right) rOvary respectively. The outgrowth expressed GFP when GFP-ESC-derived PGCLCs were used for rOvaries, indicating the proliferative outgrowths originated from GFP-ESCs. Bar scale, 200 µm. (D) Genotyping of proliferative outgrowths sampled from different 6 BVSC-iPSC-derived rOvaries. Arrowheads indicate amplified fragments targeting Blimp1-Venus, Stella-ECFP and endogenous Prdm14, respectively. All 6 proliferative outgrowths carried Blimp1-Venus and Stella-ECFP reporters, indicating the outgrowths were derived from BVSC-iPSCs. (E) BVSC-iPSC-derived oocytes with cells on the inner side of zona pellucida, harvested at day 45 of the culture. The parts in white square (left) were enlarged to right images. While Stella-ECFP was detected in the ooplasm and polar body, the contaminating cells were negative for Stella-ECFP. Bar scale, 50 µm. (F) A BVSC-iPSC-derived 2-cell embryo with cells on the inner side of zona pellucida, harvested 1 day after chemical activation of the oocyte at day 45 of the culture. The part in a black square (left top) was enlarged to an image (right top). The asterisk indicates cells on the inner side of zona pellucida. Arrow heads indicate branched zona pellucida. Bar scale, 20 µm.

**Figure S4.**
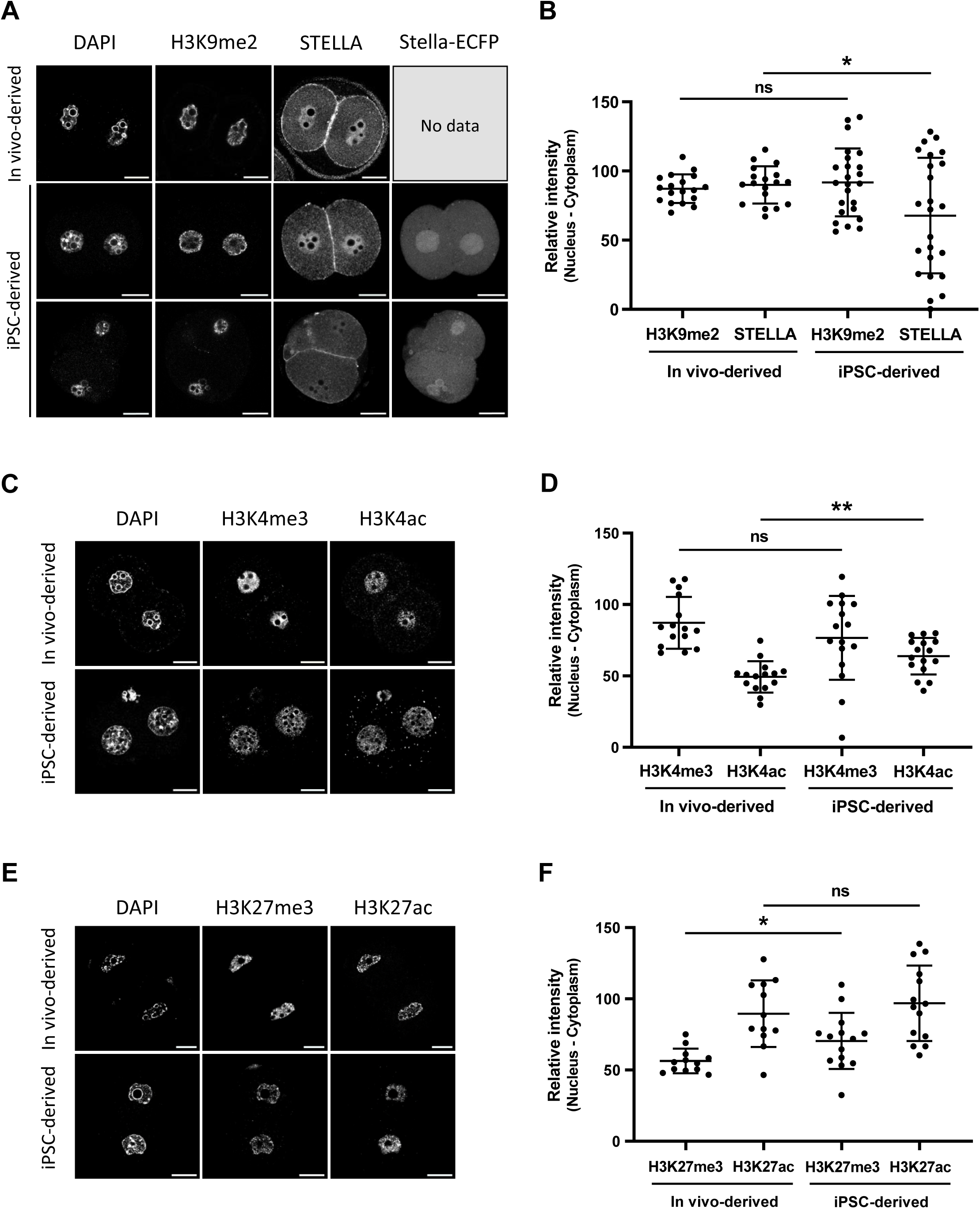
Immunostaining analysis of histone modification and STELLA in parthenogenetic 2-cell embryos. (A) Representative staining of H3K9me2 and STELLA in PA 2-cell embryos. Expression of Stella-ECFP was also shown in iPSC-derived embryos. (B) Quantitative data of H3K9me2 and STELLA staining in PA 2-cell embryos. n = 18 (*in vivo*-derived) and 24 (iPSC-derived). (C) Representative staining of H3K4me3 and H3K4ac in PA 2-cell embryos. (D) Quantitative data of H3K4me3 and H3K4ac staining in PA 2-cell embryos. n = 15 (*in vivo*-derived) and 16 (iPSC-derived). (E) Representative staining of H3K27me3 and H3K27ac in PA 2-cell embryos. (F) Quantitative data of H3K27me3 and H3K27ac staining in PA 2-cell embryos. n = 12 (*in vivo*-derived) and 14 (iPSC-derived). Bar scale, 20 µm. * P < 0.05; ** P < 0.01; ns, non-significant.

**Figure S5.**
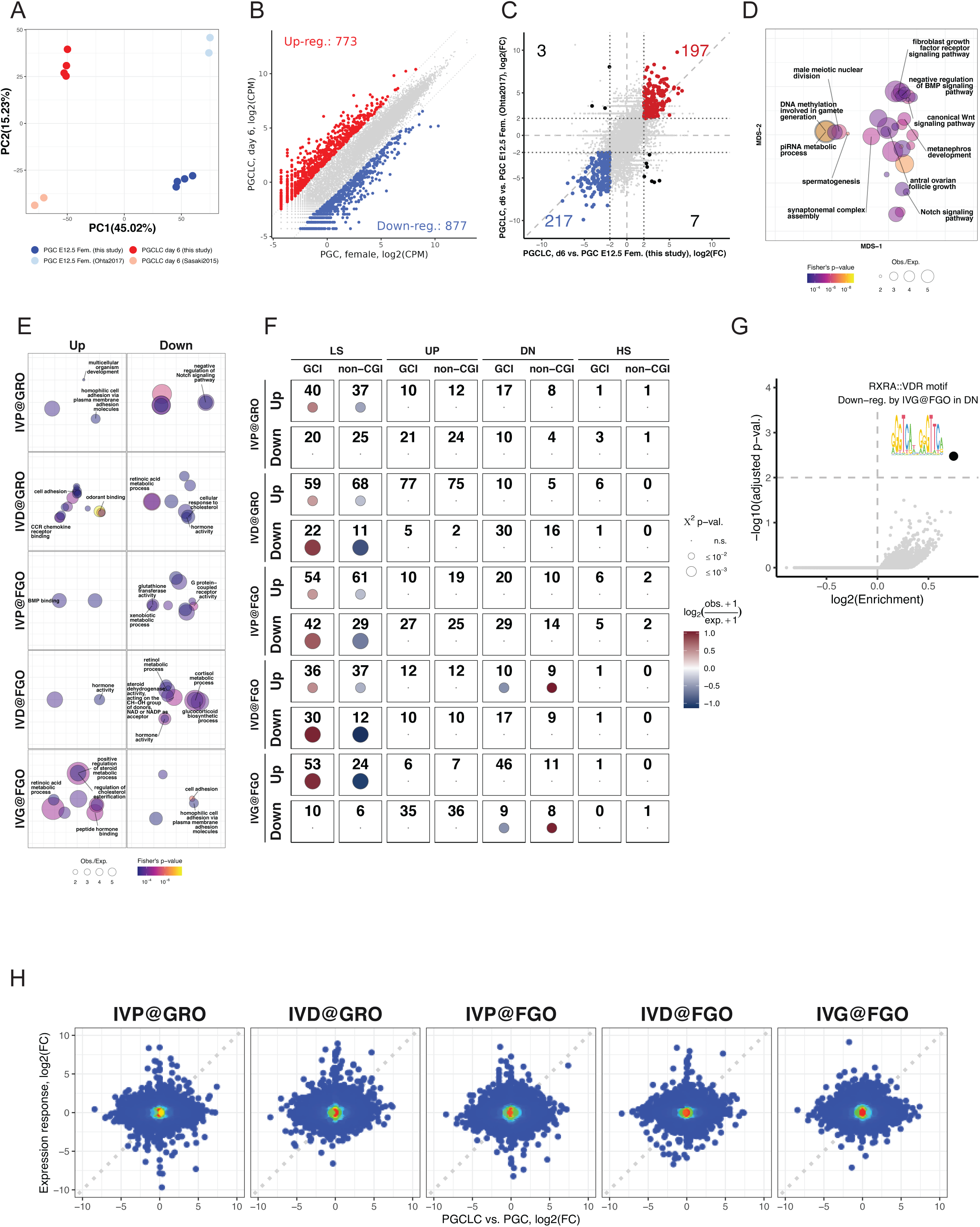
Promoter features of affected genes and comparison of genes differentially expressed in PGCLCs compared to PGCs. (A) Principal Component Analysis (PCA) of bulk RNA-Seq samples from this study for d6 PGCLC (red), PGC in E12.5 female (blue) and published RNA-Seq datasets for d6 PGCLC (pink) (Ohta et al., 2017) and PGC in E12.5 female (light blue) (Sasaki et al., 2015). PCA was performed on relative expression calculated for the dataset from this study and datasets from (Ohta *et al*., 2017) and (Sasaki *et al*., 2015) separately. (B) Scatter plot showing expression of genes in E12.5 PGCs versus d6 PGCLCs and numbers of differentially expressed genes (with *FDR* ≤ 5% and |*log*_2_(*Fold* – *change*)| ≥ 2). (C) Scatter plots illustrating correlation between expression differences between d6 PGCLC and E12.5 female PGC profiled in this study and by authors of (Sasaki *et al*., 2015) and (Ohta *et al*., 2017). Numbers of genes commonly up- and down-regulated genes in the two studies are marked in red and blue respectively (with *FDR* ≤ 5% and |*log*_2_(*Fold* – *change*)| ≥ 2). Numbers of genes with conflicting results between two studies are marked in black. (D) Gene ontology enrichment (GO) analysis for down-regulated genes in PGCLC compared to PGC (related to Table S3). Bubbles representing GO terms are scaled according to enrichments, colored according to statistical significance and positioned relative to each other to reflect similarities between significantly affected genes with corresponding GO terms (see Methods). (E) GO enrichment analysis for genes with statistically significant expression response to *in vitro* development (related to Table S6). (F) Results of *χ*^2^-tests and enrichments of genes controlled by CpG island (CGI) promoters and non-CpG island promoters (non-CGI). Enrichments with *χ*^2^-test p-value bigger than 1% are considered statistically not significant and displayed as dots. (G) Scatter plot illustrating enrichments (X-axis, log2 scale) and statistical significance (Y-axis, -log10 (adjusted p-value)) of transcription factor motifs in promoters of genes which are up- or down-regulated by each stage of *in vitro* development and display particular expression dynamics from GRO to FGO *in vivo* (Figures 5C and 5E). (H) Scatter plots showing expression differences between E12.5 PGCs and d6 PGCLCs versus expression responses to each stage of *in vitro* development.

**Figure S6.**
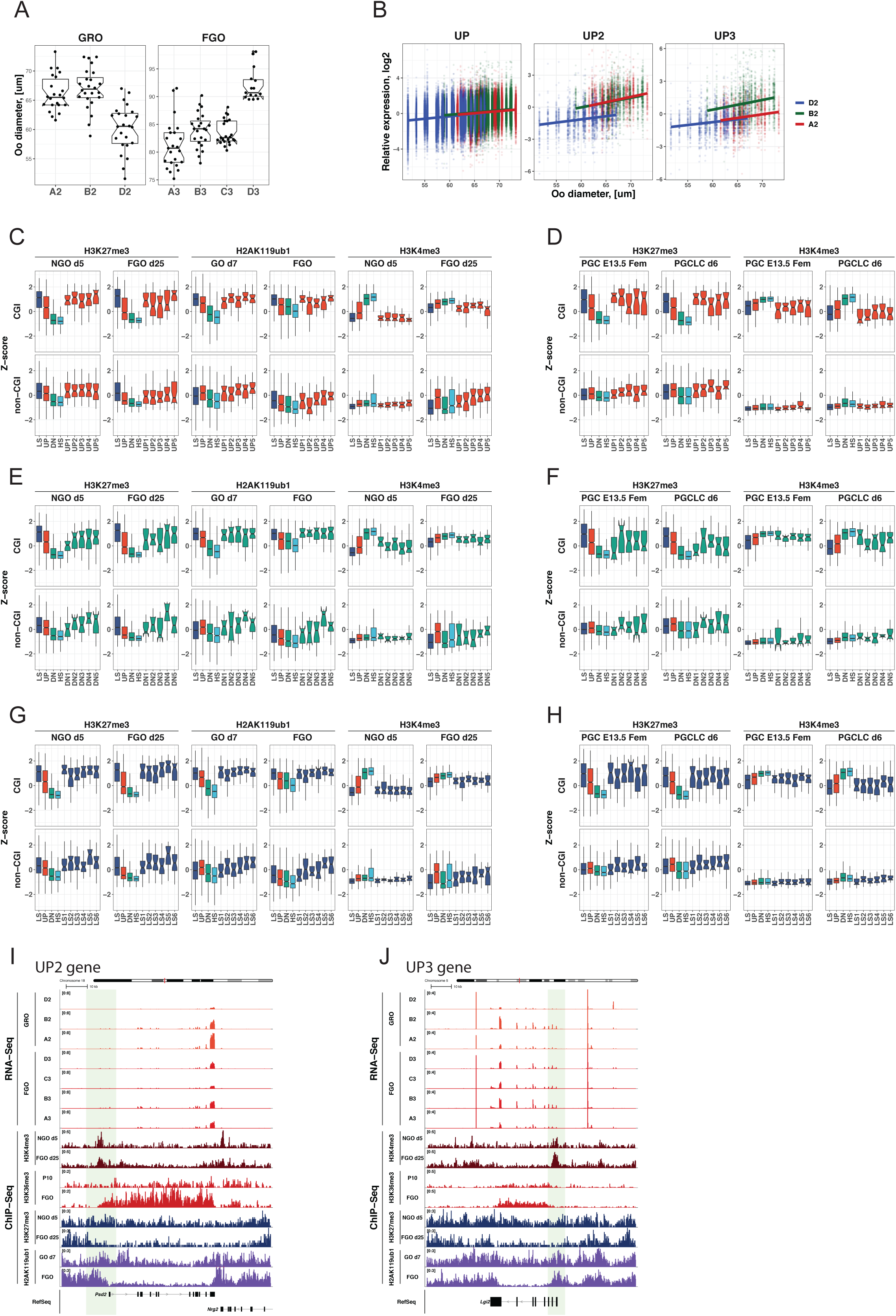
Investigation of chromatin features of promoters of genes affected by *in vitro* PGCLC and oogenesis procedure. (A) Boxplots illustrating differences in sizes of oocytes belonging to different cohorts of GRO (A2, B2, D2) and FGO (A3, B3, C3 and D3). (B) Investigation of possible confounding effect of oocyte size differences on results of differential expression analysis in GRO. Each point represents relative expression of a gene (Y-axis) in a particular GRO with corresponding size in X-axis. Left panel shows relative expression of all genes showing up-regulation dynamics (UP group) in GROs of different cohorts (A2 in red, B2 in green, D2 in blue), middle and rights panels represent relative expression of genes belonging to UP2 and UP3 clusters shown in Figure 6B respectively. (C) Enrichments (scaled and centered to Z-scores) of H3K27me3 in non-growing oocytes and FGO (data from (Hanna et al., 2018)), H2AK119Ub1 in growing oocytes at day 7 and FGO (data from (Mei et al., 2021)) and H3K4me3 in non-growing oocytes and FGO (data from (Hanna *et al*., 2018)) at CGI and non-CGI promoters (+/- 1.5 kb) of genes with specific expression dynamics *in vivo* (LS, UP, DN, HS groups, see Figure 5C) and affected genes in UP cluster from Figure 6B. (D) Enrichments (scaled and centered to Z-scores) of H3K27me3 and H3K4me3 marks in PGC E13.5 (data from (Kawabata et al., 2019)) and d6 PGCLC (data from (Kurimoto et al., 2015)) at CGI and non-CGI promoters (+/- 1.5 kb) of genes with specific expression dynamics *in vivo* (LS, UP, DN, HS groups, see Figure 5C) and affected genes in UP cluster from Figure 6B. (E) Analogous to (C) but for affected genes in DN cluster from Figure 6B. (F) Analogous to (D) but for affected genes in DN cluster from Figure 6B. (G) Analogous to (C) but for affected genes in LS cluster from Figure 6B. (H) Analogous to (D) but for affected genes in LS cluster from Figure 6B. (I) Genomic snapshot of a representative gene *Psd2* belonging to the UP2 cluster (see Figure 6B) and distribution of several chromatin marks. (J) Genomic snapshot of a representative gene *Lgi2* belonging to the UP3 cluster (see Figure 6B) and distribution of several chromatin marks.

